# Partitioning genetic effects on birth weight at classical human leukocyte antigen loci into indirect maternal and direct fetal components using structural equation modelling

**DOI:** 10.1101/2022.08.23.505053

**Authors:** Geng Wang, Nicole M Warrington, David M Evans

## Abstract

Birth weight (BW), as a proxy for intrauterine growth, is influenced by both fetal and maternal genetic factors. Single nucleotide polymorphisms in the human leukocyte antigen (HLA) region in both maternal and fetal genomes have been robustly associated with BW in previous genetic association studies suggesting the involvement of classical HLA alleles in BW etiology. However, no study to date has partitioned the association between BW and classical HLA alleles into maternal and fetal components. We used structural equation modelling (SEM) to estimate the indirect maternal (i.e. via the intrauterine environment) and direct fetal effects of classical HLA alleles on BW. Our SEM leverages the data structure of the UK Biobank (UKB), which includes participants’ own BW and/or the BW of their firstborn child (in the case of UKB females). We show via simulation that our model yields asymptotically unbiased estimates of the maternal and fetal allelic effects on BW and appropriate type I error rates, in contrast to simple regression models that estimate unconditioned maternal and fetal effects. Asymptotic power calculations show that we have sufficient power to detect moderate-sized maternal or fetal allelic effects (standardized effect size ≥ 0.01) of common HLA alleles on BW in the UKB. Applying our SEM to imputed classical HLA alleles and own and offspring BW of ∼270,000 participants from the UKB replicated the previously reported association at the *HLA-C* locus (*C*04:01*, P = 2.13×10^−7^, *C*05:01, P=* 6.91×10^−5^, *C*03:03*, P= 4.53×10^−3^, respectively) and revealed strong evidence for maternal (*HLA-A*03:01*, P = 7.90×10^−8^; *B*35:01*, P = 7.78×10^−5^; *B*39:06*, P = 8.49 ×10^−5^) and fetal allelic effects (*HLA-B*39:06*, P = 4.03×10^−4^) of non-*HLA-C* alleles on BW. These novel allelic associations between BW and classical HLA alleles provide insight into the immunogenetics of fetal growth *in utero*.

## Introduction

Birth weight (BW) is widely used as a cheap but imperfect measure of intrauterine growth and development^1,2^. Both high and low BW infants are at increased risk of short- and longterm morbidities. Low BW infants are at increased risk of neonatal mortality^3^ and cardiometabolic diseases in later life^4^, which in the latter’s case, might reflect “developmental programming” as a consequence of an adverse intrauterine environment ^5^. In contrast, high BW infants (fetal macrosomia) are at increased risk of injury during birth (e.g. delivery shoulder dystocia, obstructive labour)^6^ and type 2 diabetes in later life^7^.

BW is a complex trait influenced by both maternal and fetal genetics and environmental factors^8^. Large-scale genome-wide association studies (GWAS)^8-12^ have identified over 200 loci associated with BW, including single nucleotide polymorphisms (SNPs) in the human leukocyte antigen (HLA, also known as major histocompatibility complex[MHC]) region on the short arm of chromosome 6^8,12^. The classical HLA antigens can be divided into HLA class I (*HLA-A, B*, and *C*) and class II (*HLA-DP, -DQ*, and *-DR* loci)^13^. This area of the genome exhibits extensive and complex patterns of linkage disequilibrium and contains a set of highly polymorphic genes that encode cell surface antigens that play a key role in specific immunity, allograft rejection and the development of many immune-mediated and autoimmune diseases (reviewed in^14^). These results are interesting, given that epidemiological and immunological studies have suggested that both maternal and fetal immune systems play important roles in fetal growth and development ^13,15-17^

One complication with interpreting the results of genetic association studies of perinatal traits, is that it is often unclear whether specific genetic associations reflect an effect of the fetal genome, an effect of the maternal genome, or some mixture of both (Evans et al. 2019).

Insight into this question can sometimes be obtained by performing conditional genetic association analyses (i.e. where the effect of fetal genotype is conditioned on maternal genotype and vice versa) and/or haplotype analyses of genotyped mother-offspring pairs^12^. A complication, however, is that worldwide there is a paucity of cohorts with large numbers of genotyped mother-offspring pairs meaning that these sorts of analyses are often underpowered ^18^. In order to address this issue, Warrington et al. (2018) developed a structural equation model that was capable of partitioning genetic effects on BW into maternal and fetal components using the large UK Biobank (UKB) resource^19^. In the UKB, genotyped participants report their own BW, and the BW of their first child (females only)^20^. Wu et al. (2021) further extended this statistical framework to estimate parental and fetal genetic effects using summary results statistics from GWAS conducted on own and offspring BW whilst simultaneously accounting for sample overlap and one generation of assortative mating^21^. However, these models were derived for bi-allelic variants (mostly SNPs), whereas classical HLA genes are multi-allelic. Thus, an extension of the model is required to estimate maternal and fetal effects at multi-allelic classical HLA genes.

In this manuscript, we develop a novel structural equation model to estimate the maternal and fetal genetic effects of imputed HLA alleles on offspring outcomes. We conducted a simulation study to assess the bias and the type I error rate of our model and performed asymptotic power calculations to estimate the power to detect the allelic effects and partition them into maternal and fetal allelic effects. We subsequently apply our structural equation modelling (SEM) to data from the UKB to estimate the maternal and fetal genetic effects of classical HLA alleles on BW.

## Methods

### Structural Equation Modelling

Structural equation modelling (SEM), with its ability to model the relationship between latent and observed variables, provides a natural framework for investigating the association between maternal and fetal genotypes and perinatal phenotypes^22^. We illustrate the SEM we developed to estimate maternal and fetal genetic effects on BW at classical HLA alleles in the form of a path diagram in Figure 1. To simplify explication, we assume the locus contains only four hypothetical alleles (here *HLA-X0, HLA-X1, HLA-X2, HLA-X3*, Fig. 1) in the fictitious *HLA-X* gene (we use “*HLA-X*” to avoid confusion with real HLA genes), although the model can be generalized, at least in theory, to an arbitrary number of alleles. We set the most common allele, here the *HLA-X0* allele, as the “baseline”, and is hence not included in the SEM model to avoid collinearity. The model contains both observed variables (represented by squares in the path diagram) and latent variables (represented by circles in the path diagram). The two observed phenotypic variables are the self-reported BW of the individual (BW) and the self-reported BW of their first offspring in the case of women in the UKB (BW_O_). The three observed genetic variables are *HLA-X1*_*M*_, *HLA-X2*_*M*_, and *HLA-X3*_*M*_, which represent the number of copies of each of these alleles that each individual carries (the “M” subscript referring to the mothers in the SEM in Figure 1). The rest of the variables are modelled as latent variables. We estimate the maternal and fetal effects of all the alleles at one locus simultaneously and the covariances across all the alleles. However, in a more realistic scenario, not every individual reports both phenotypes (i.e., they report only BW or BW_O_). In order to include the maximum amount of data to optimize power, we also simultaneously modelled individuals who failed to report either their own or their offspring’s BW (Supplementary Figure 1). The model was written and fit using the OpenMx package^23^ (version 2.19.8) in R (version 3.6.3; the R script for conducting analysis using our model is provided in the Supplementary Note).

**Figure 1.**
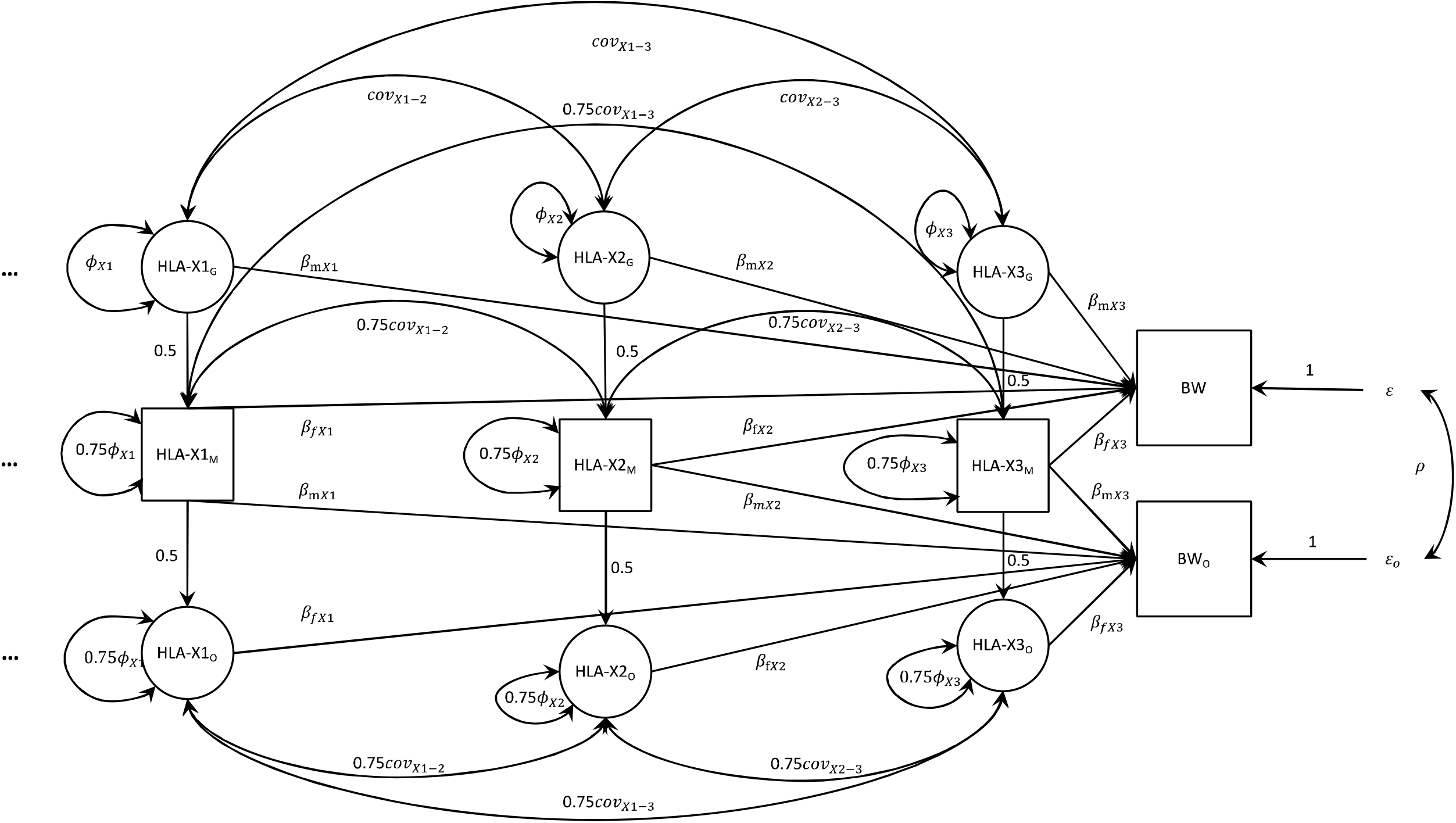
Structural equation model used for the analysis of multi-allelic HLA markers and birth weight Latent variables are represented by circles, observed variables by square boxes. Causal paths are indicated by unidirectional arrows, bidirectional arrows represent covariances. In order to illustrate the model we use the fictional example of the *HLA-X* gene. We assume there are only four alleles in the *HLA-X* gene (here *HLA-X0, HLA-X1, HLA-X2, HLA-X3*), although the model is generalizable to an arbitrary number of alleles (represented by the ellipsis “…” on the left-hand side of the path model). The *HLA-X0* allele is modelled as the “baseline” allele (so all effects are modelled as displacements from this baseline genotype), and hence is not shown in the diagram. The *HLA-X1, HLA-X2*, and *HLA-X3* variables represent the number of copies of each of these alleles that each individual carries. The “G”, “M”, and “O” subscripts index genotypes in the grandmaternal (latent), maternal (observed), and offspring (latent) generation respectively. The “X1”, “X2”, and “X3” subscripts index the corresponding alleles. The “m” and “f” subscripts of the path coefficients represent maternal and fetal allelic effects, respectively. The five observed variables, displayed in squares, in the analysis are 1) the self-reported birth weight of the individual (BW) in the UKB, 2) the self-reported BW of their first offspring (in the case of women only) in the UK Biobank (BW_O_), and 3) the number of copies of the *HLA-X1* (*HLA-X1*_*M*_), *HLA-X2* (*HLA-X2*_*M*_), and *HLA-X3* (*HLA-X3*_*M*_) alleles. The latent variables, displayed in circles, in the analysis are 1) the number of copies of the relevant allele carried by the individual’s mother (i.e. *HLA-X1*_*G*_, *HLA-X2*_*G*_, and *HLA-X3*_*G*_) and 2) the number of copies of the relevant allele carried by the individual’s offspring (*HLA-X1*_*O*_, *HLA-X2*_*O*_, and *HLA-X3*_*O*_). The variance of the allele counts in the grandmaternal (*HLA-X1*_*G*_, *HLA-X2*_*G*_, *HLA-X3*_*G*_), maternal (*HLA-X1*_*M*_, *HLA-X2*_*M*_, *HLA-X3*_*M*_) and offspring (*HLA-X1*_*O*_, *HLA-X2*_*O*_, *HLA-X3*_*O*_) generations are estimated as Φ_X1_, Φ_X2_ and Φ_X3_, respectively, and assumed not to change across generations (i.e., variance (*HLA-X*_G_) = Φ, variance (*HLA-X*_M_) = 0.75Φ + 0.25Φ = Φ, and variance (*HLA-X*_O_) = 0.75Φ + 0.25Φ = Φ). The covariances between the different allele counts are also estimated (e.g. cov_X1-2_, cov_X1-3_, cov_X2-3_) and assumed not to vary across generations (e.g., covariance (*HLA-X1*_*G*_, *HLA-X2*_*G*_) = cov_X1-2_, covariance (*HLA-X1*_*M*_, *HLA-X2*_*M*_) = 0.75 cov_X1-2_ + 0.25 cov_X1-2_ = cov_X1-2_, and covariance (*HLA-X1*_*O*_, *HLA-X2*_*O*_) = 0.75 cov_X1-2_ + 0.25 cov_X1-2_ = cov_X1-2_).. The β_*mX*1_, β_*fX*1_, β_*mX*2_, β_*fX*2_, β_*mX*3_ and β_*fX*3_ are path coefficients that quantify the maternal and fetal allelic effects of each allele on BW, respectively. The residual error terms for the BW of the individual and their offspring are represented by □ and □_o_ respectively, and the variance of both of these terms is estimated in the SEM. The estimated covariance between residual genetic and environmental sources of variation on BW is represented by ρ.

### Simulations to Assess Bias and Type I Error

We performed simulations to investigate the accuracy of our SEM to estimate both maternal and fetal effects on BW. For each scenario, we generated 1,000 replicates where we analyzed 80,000 genotyped individuals who had simulated phenotypic data for their own BW and the BW of their offspring. For each replicate, we generated HLA alleles for the three generations of individuals at a single locus (i.e. even though only the HLA alleles of the individual in the middle generation was modelled as an observed variable in the analysis). The variances of each allele was standardized to unit variance. For simplicity, we only simulated one locus and assumed the absence of linkage disequilibrium between the locus and other classical HLA genes in the MHC region. We generated the individual’s own BW variable (BW) for each individual *j* using the following equation (Eq. 1):

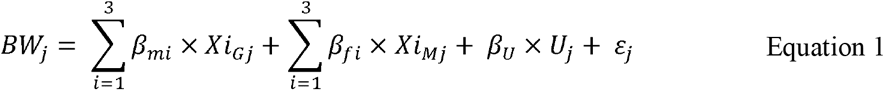

where *β*_*m,i*_ indicates the additive effect of an individual’s mother’s allele *HLA-Xi* (i = 1…3) on their offspring’s BW relative to an individual whose mother is homozygous for the “baseline” allele *HLA-X0* (i.e. the indirect maternal effect), and *Xi*_*Gj*_ represents the number of copies of the *HLA-Xi* allele in individual *j*’s mother (the subscript G is used here because this is the grandmaternal generation and to make the notation in our simulations correspond with that in the path diagram in Figure 1). *β*_*fi*_ is the effect of an individual’s own allele *HLA-Xi* on their own BW (i.e. the direct fetal effect). *Xi*_*Mj*_ represents the number of copies of the *HLA-Xi* allele in individual *j* (the subscript M is used here because this is the maternal generation). *U*_*j*_ is a standard normal random variable that denotes all residual shared genetic and environmental components that influence own and offspring birthweight across the generations for individual *j. β*_*U*_ is the effect of *U*_*j*_ on the individual’s own BW, and *ε*_*j*_ is a random normal variable with mean zero and variance selected to ensure that BW has unit variance asymptotically.

Similarly, offspring BW (BW_O_), for each individual *j*, was generated using the following equation (Eq. 2):

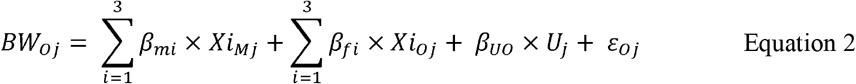

where *Xio*_*j*_ is a latent variable representing the number of copies of *HLA-Xi* allele (i = 1…3) in the offspring *j, β*_*Uo*_ is the effect of residual genetic and environmental influences *U* on offspring BW, *ε*_*0*_ is a random normal variable with mean zero and variance selected to ensure that *BW*_*o*_ has unit variance asymptotically and the other variables are defined as above.

We simulated the classical alleles *HLA-X0, HLA-X1, HLA-X2*, and *HLA-X3* to have frequencies = 0.7, 0.1, 0.1, and 0.1 respectively in 80,000 mother-offspring pairs. The corresponding fetal effects (*β*_*fi*_) were set to *β*_*f1*_ = 0, *β*_*f2*_ = 0.014, *β*_*f3*_ = 0.02 (Table S1) and the maternal effects (*β*_*mi*_) of these allele were assigned as *β*_*m1*_ = 0, *β*_*m*2_ = 0.01, *β*_*m*3_ = 0.014 (Table S1). We also simulated the situation where maternal and fetal effects were in opposite directions (i.e. *β*_*mi*_ = 0, -0.01, and -0.014). The allele *HLA-X0* was set to the baseline allele in each scenario.

For each simulation, we then fit both simple linear regression and SEM models to the data. The linear model regressed either offspring BW (*BW*_*O*_) or own BW (*BW*) on own classical HLA alleles (*Xi*_*Mj*_; which is similar to estimating a maternal or fetal allelic effect without properly conditioning on fetal or maternal allele). We fit the SEM displayed in Figure 1 to estimate the independent maternal and fetal effects of HLA alleles simultaneously. Bias and type I error rates were calculated in each scenario. Bias was defined as the mean difference between the estimated allelic effects and the true simulated allelic effect sizes across all simulations and was calculated for both maternal and fetal allelic effects. Type I error rate was calculated as the proportion of tests that reached P < 0.05 under the null hypothesis when the true allelic effect of interest was zero. Monte Carlo standard errors for bias and type I error rate were calculated^24^.

### Asymptotic Power Calculations

We also performed asymptotic power calculations using the OpenMx package (version 2.19.8 in R (version 3.6.3). In brief, asymptotic covariance matrices were generated assuming the same underlying values for the parameters of the model as in the simulations above. We first estimated the parameter values using the full model, which was fitted to the covariance matrices, and checked that this provided a perfect fit to the data. Second, we constrained the parameter(s) of interest to zero (i.e. the maternal allelic effect, or the fetal allelic effect or both of these parameters simultaneously) in a reduced model fitted to the same covariance matrices. The difference in *χ*^2^ between the full and reduced models is equal to the non-centrality parameter of the test for association under the alternative hypothesis with the degrees of freedom equal to the number of constrained parameter(s) in the reduced models (i.e. one or two). Power was then estimated as the area under the curve of the rejection region (i.e. under a non-central *χ*^2^ distribution to the right of the significance threshold of interest [α= 0.05], Eq. 3).

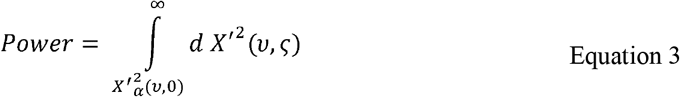

where 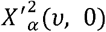 is the quantile of the 100 × (1□−□ *α*) percentage point of the central *χ*^2^ distribution with a type I error rate of a and degrees of freedom *ν*, and *ς* is the non-centrality parameter. For further details of the procedure see the description by Moen et al.^18^

In addition to power calculations performed assuming complete data, we also calculated power when individuals only reported their own or their offspring’s BW. We assumed 100,000 individuals who only reported their own BW, 80,000 mothers who only reported the birthweight of their firstborn and 80,000 individuals who reported both (i.e. similar numbers of individuals as in the UKB). We fit an SEM that included additional structures for individuals who only reported their own or their offspring’s birthweight (Supplementary Figure 1).

### Application of Model to BW Data in the UK Biobank

The UK Biobank (UKB) Study is a study of over 500,000 volunteers (with 5.45% response rate of those invited ^25^), recruited from across the UK at age 40-69 years between 2006 and 2010, with a broad range of health-related information and genome-wide genetic data ^26^. The UKB samples were genotyped using one of two different Affymetrix arrays (the UK BiLEVE Axiom array or the UK Biobank Axiom array). Genotyping, quality control, and imputation were performed centrally by the UKB^26^. Genotype and phenotype data are available upon application to the UKB (http://www.ukbiobank.ac.uk/); the results of this study were based on data accessed using approved applications ID 53641. For our analysis, we used HLA data imputed using HLA*IMP02^27^ by the UKB. We used the “best guess” genotypes (i.e., 0, 1, 2 for each HLA allele per individual) derived from posterior allelic probabilities (Q statistics from HLA*IMP02 imputation) and set all genotypes with a Q below 0.7 to missing. The UKB has approval from the North-West Multi-Centre Research Ethics Committee, which covers the United Kingdom and all participants provided written informed consent.

We defined a subset of participants of “European” origin by conducting an ancestry informative principal components (PC) analysis using participants from Phase 3 of the 1000 Genomes project ^28^ as a reference for ancestry. Directly genotyped data were used for ancestry determination. The UKB participants were then projected into this PC space according to the SNP loadings generated from a 1000 Genomes PC analysis using FlashPCA2^29^. PC1, PC2 and PC5 resolved the British/European (GBR/CEU [i.e. British in England and Scotland/Western European Ancestry]) cluster efficiently and were hence used in subsequent clustering. The UKB participants’ ancestry was classified using an Expectation-Maximization Clustering (EMC) algorithm (https://CRAN.R-project.org/package=EMC) centred on the 26 different 1000 genomes populations. After comparing how well the EMC clustering model fit the data using the chosen PCs and by varying the numbers of predefined clusters (1-50 clusters), 12 clusters showed an optimal balance between improved model fit and resolution. Those UKB participants clustering with the GBR/CEU clusters were classified as having “white British” ancestry.

In UKB, there are 280,065 participants who reported their own BW, and 216,771 mothers who reported the BW of their first born offspring^20^ at any one of three occasions (baseline or follow-up). If participants had an own/offspring BW reported at baseline, then this measure was used for analysis; otherwise, we used BW reported at either follow-up one or two (in this order). For those participants who reported their BW on more than one occasion, we excluded BW measurements if the difference between two reported BWs were > 1kg. UKB does not record the gestational age of mothers, so we further excluded the measurements likely to be either pre-term or postterm births. For self-reported BW, we excluded the BWs of individuals < 2.5 kg or > 4.5 kg. For mother-reported offspring BW, we excluded the offspring BW < 2.27 kg (5 pounds) or > 4.54 kg (10 pounds; the weights of firstborn were recorded to the nearest whole pound in UKB). We excluded those UKB participants who self-reported being part of multiple births. Only genotyped individuals that passed quality control by UKB^26^ were included in further analyses.

Related individuals were identified using KING software^30^ (we define unrelated as pairs of individuals who were less related than 3^rd^-degree relatives as defined by the KING software, i.e. kinship coefficient 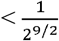). One individual from each related pair was then excluded from analyses to maximize the number of independent individuals reporting BW. After cleaning, 105,121 unrelated European individuals who only reported their own BW, 82,445 mothers who reported only the BW of their firstborn, and 85,757 mothers who reported both were retained for subsequent analyses (Supplementary Figure S2 and S3). Z-scores of individuals’ own BW and the BW of their first child were generated after adjusting for both the top 40 GWAS-derived principal components and for sex in the case of own BW (sex was not reported for the offspring).

We fit our SEM to UKB self-reported BW, offspring BW, and allelic status at 11 classical imputed HLA loci (*HLA-A, HLA-B, HLA-C, HLA-DRB1, HLA-DRB3, HLA-DRB4, HLA-DRB5, HLA-DQB1, HLA-DQA1, HLA-DPB1*, and *HLA-DPA1*) using the OpenMx package in R (version 3.6.3). HLA alleles with a frequency lower than 0.5% were grouped together in the analyses (only when the total frequency of grouped alleles was over 0.5%).

In addition to one degree of freedom tests where we evaluated the significance of the maternal and fetal components individually, we also compared the full model with a constrained model in which we fixed the maternal and fetal effects of the allele of interest to zero (i.e. a two degree of freedom test). P values were calculated by comparing the full and constrained models using the difference in minus twice the log-likelihood chi-square with degrees of freedom equal to the difference in number of free parameters between the models (i.e. two). These two degree of freedom tests therefore test whether there is evidence for any effect of the allele on BW (i.e. regardless of whether it is operating through the maternal or fetal genomes or both) at the locus of interest.

The top BW-associated SNPs in the MHC region reported by Warrington et al. (2019) ^8^ were rs9366778 (near *HLA-C*) and rs6911024 (in the 5’ UTR of *MICA*[MHC class I polypeptide– related sequence A]). To examine the relationship between these SNPs, the classical HLA alleles in the region and BW, we conducted conditional analyses whereby we augmented our SEM by including these SNPs as observed variables in the model. We modelled the contribution of grandmaternal and offspring genotypes at the same SNPs as latent variables (Supplementary Figure S4). This model therefore effectively conditions the HLA results on these SNPs and allows us to examine whether classical HLA alleles are associated with birthweight independent of the SNP associations. The model also accounts for any linkage disequilibrium between the SNPs and HLA alleles, as these are included as (estimated) covariances in the SEM.

## Results

### Bias and Type I Error Rate

Figure 2 and Table S1 show the bias in estimating maternal and fetal allelic effects of the simulated *HLA-X* gene using linear regression or the SEM (results are shown for the *HLA-X1* allele only, but other alleles show similar patterns). Our results suggest the SEM yields asymptotically unbiased estimates of maternal and fetal effects in contrast to ordinary least squares regression, which does not explicitly partition allelic effects into maternal and fetal components. Likewise, whilst the SEM produced well calibrated type I error rates (≤5%) for tests of the estimated effects when the true effect size was set to zero, the type I error rates for linear regression could be inflated e.g. when the true maternal effect was zero but fetal effects were present (or vice versa) (Table S2).

**Figure 2.**
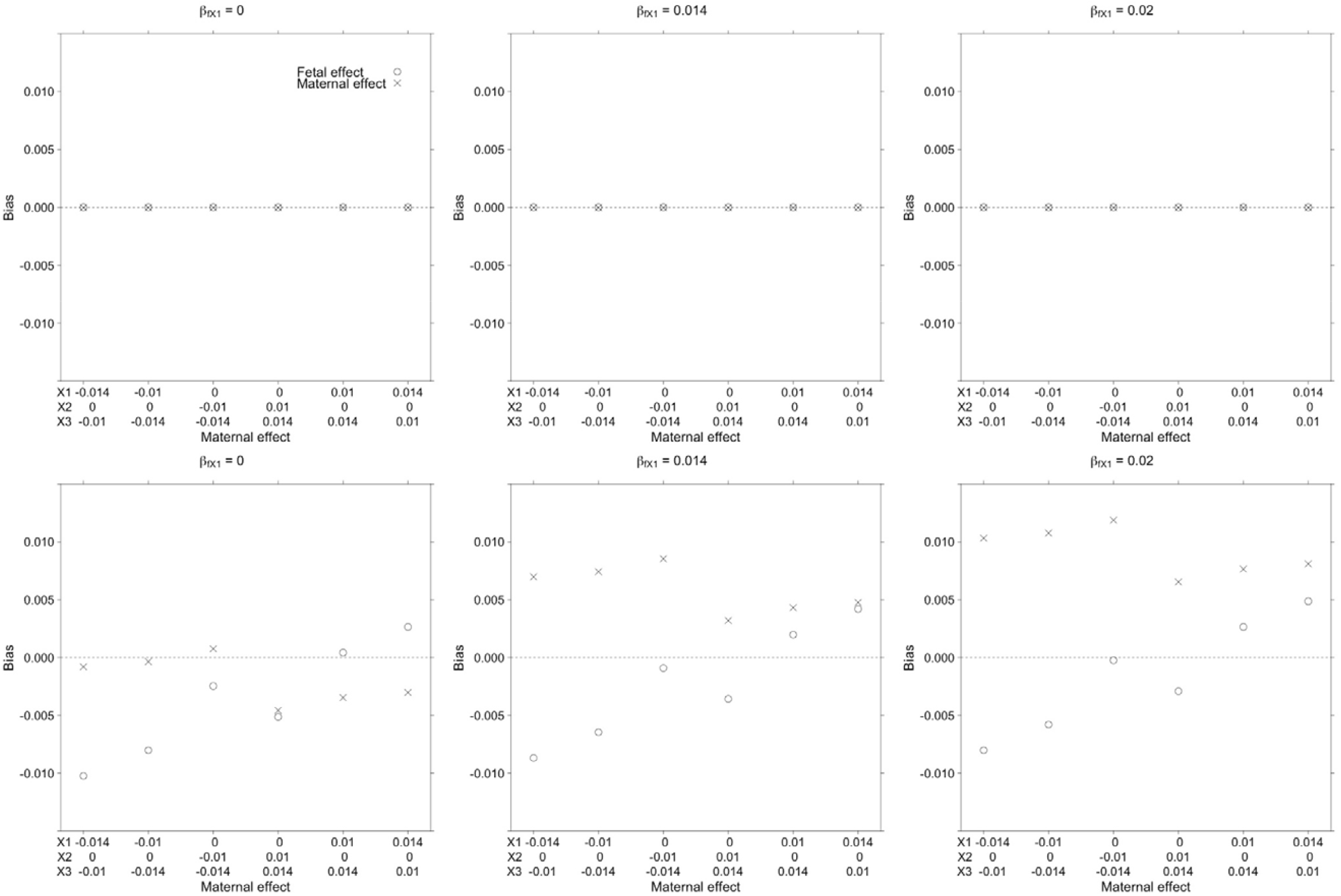
Simulation results showing bias in estimates of the maternal and fetal effect of the HLA-X1 allele obtained using structural equation modelling (upper row) or ordinary least squares linear regression (lower row). The true fetal effect of *HLA-X1* varies between the six panels (β_fX1_ = 0 left panels; β_fX1_ = 0.014 middle panels; β_fX1_ = 0.02 right panels - the fetal effects of other HLA-X alleles for each panel are listed in Supplementary Table S1). The effect of the maternal alleles (β_mX_1, β_mX2_, β_mX3_) varies across the x-axis. Across all conditions, the SEM returned unbiased estimates of the maternal and fetal effect size for *HLA-X1* whereas ordinary least squares regression resulted in biased effect estimates. Similar patterns of results were observed for *HLA-X* alleles other than *HLA-X1* (results not shown).

### Asymptotic Power Calculation

Figure 3 and Table S3 present asymptotic power to detect the effect of classical HLA alleles (i.e. a two degree of freedom test) and subsequently partition the allelic effect into maternal and fetal components (i.e. one degree of freedom tests) when the sample size is set to 80,000 mothers who report their own and their offspring’s phenotype (α = 0.05). We have excellent power (> 80%) when the standardized effect of the allele of interest (*HLA-X1*; allele frequency = 0.1) is greater than 0.014 (the variances of all alleles have been standardized to unit variance). Even with a maternal standardized effect size as low as 0.01, the model still had > 50% power to detect the partitioned maternal effect. As expected, the two degree of freedom test had more power (to detect any allelic effect at the locus) than the one degree of freedom maternal/fetal tests for all conditions we examined. The asymptotic power calculation is consistent with the results of simulation study (Table S2). The addition of incomplete data (i.e. individuals who report only their own phenotype or alternatively their offspring’s phenotype, but not both) also increased power (Figure 3; Table S4).

**Figure 3.**
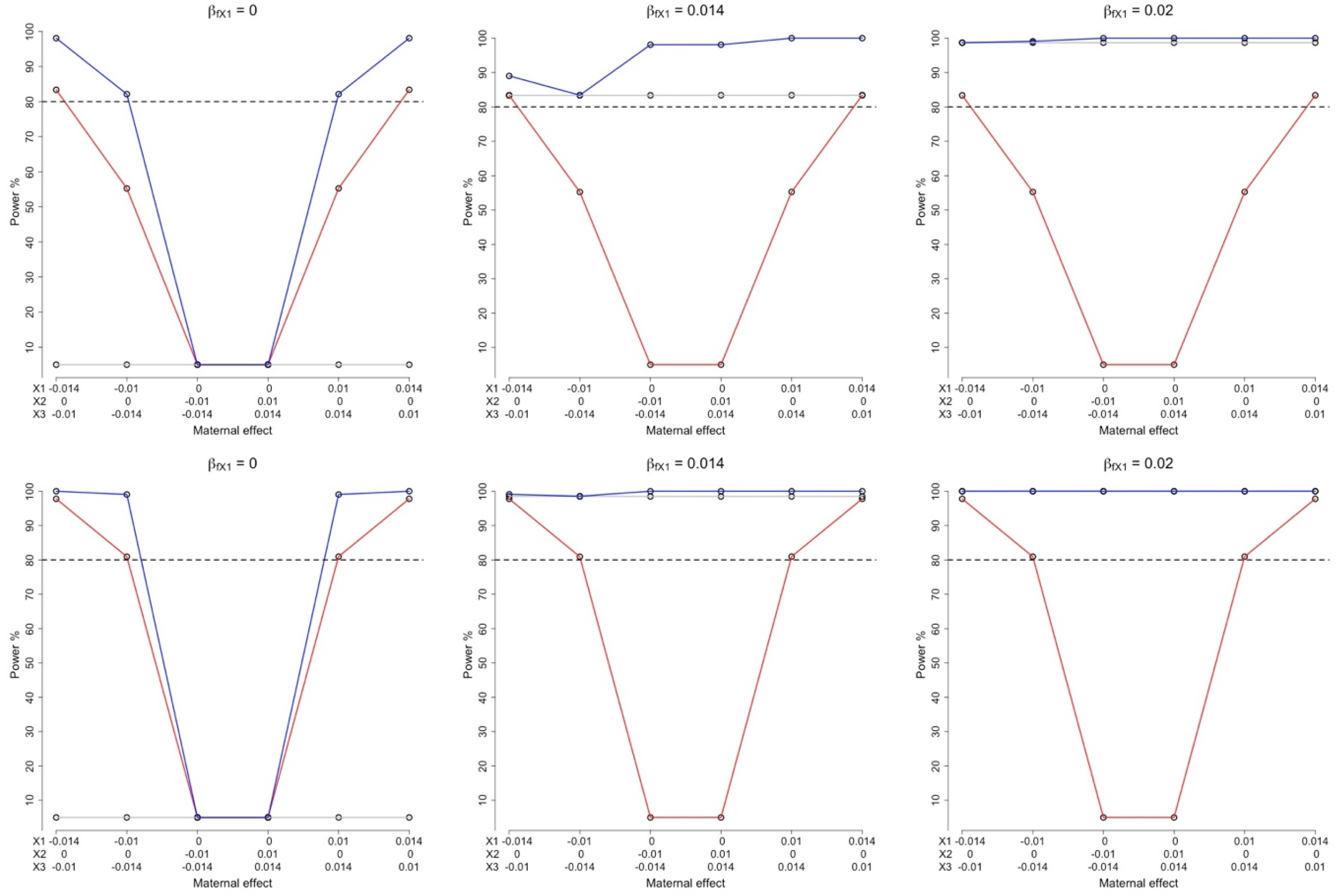
Asymptotic power of the structural equation model Power of the SEM to detect (i.e. a two degree of freedom test where both maternal and fetal components are set to zero) and partition the effects (one degree of freedom tests) of different combinations of underlying maternal and fetal effects (α = 0.05). The colours of the lines represent tests for fetal (grey), maternal (red) or both (blue) components. β_*m*_: maternal effect; β_*f*_: fetal effect. The horizontal dashed line shows the level for 80% power. The three plots on the upper row show power calculations for 80,000 samples with both own and offspring birth weight (BW). The three plots on the lower row show power calculations for 80,000 samples with both own and offspring BW, 100,000 samples with their own BW only, and 80,000 samples with offspring BW only.

### Empirical Analysis in UK Biobank

The most common alleles for each locus (which were set to the “baseline” allele in the SEM) were *HLA-A*02:01, -B*07:02, -C*07:01, -DRB1*03:01, -DRB3*99:01, -DRB4*99:01, - DRB5*99:01, -DQB1*03:01, -DQA1*05:01, -DPB1*04:01, and -DPA1*01:*03 (Supplementary Table S5). In the case of the HLA-DRB genes, whilst the *HLA-DRB1* gene is present in all individuals ubiquitously, either none or one of the functional *DRB3/4/5* genes is present in each individual. In the UKB HLA imputation, the allele “99:01” for *HLA-DRB3, - DRB4*, and *-DRB5* genes indicate the absence of a functional locus. In total across all loci, there were terms for 124 HLA alleles included in the analyses (including seven terms where rare alleles were grouped together). We did not include terms for rare alleles at four loci because the remaining frequencies were < 0.5% in total (*HLA-DPA1* and *-DRB3*) or all imputed alleles at the locus had frequencies over 0.5% (*HLA-DRB4* and *-DRB5*).

In the two degree of freedom tests, there were a total of 29 alleles that showed nominal association with BW (P < 0.05, Supplementary Table S6). The most significant alleles were *B*35:01* (P_2df_ = 2.45×10^−9^), *A*03:01*(P_2df_ = 7.39×10^−8^), *C*04:01*(P_2df_ = 2.13×10^−7^), *C*05:01*(P_2df_ = 6.91 ×10^−5^), *B*39:06*(P_2df_ = 4.45 ×10^−4^), and *DPB1*01:01* (P_2df_ =5.49×10^−4^).

Figure 4 summarizes the results of partitioning the effects at HLA loci into maternal and fetal components in the UKB dataset. A total of nineteen HLA alleles had nominally significant maternal and/or fetal effects on BW (P < 0.05; Supplementary Table S7). Twelve of the nineteen alleles had evidence for a maternal effect only, four alleles primarily had evidence for a fetal effect only, and three alleles had evidence of both. We further set a more stringent threshold for significance using Bonferroni correction for 50 tests (α = 0.05/50 = 0.001), on the rationale that the locus with the most coded alleles (HLA-B, 25 alleles) involved 50 statistical tests (i.e. one maternal and one fetal effect for each coded allele). Three of the alleles exhibited significant evidence for maternal effects with P values less than 0.001 (*A*03:01, B*35:01, B*39:06*, labelled in Figure 4), and the *HLA-B*39:06* allele exhibited a significant fetal effect at P < 0.001 (labelled in Figure 4). The allele that exhibited the most significant p-value, *A*03:01*, showed evidence for opposite maternal and fetal effects (i.e. maternal effect = -0.042 [-0.057, -0.027], P_m_ =7.90×10^−8^; fetal effect = 0.020 [0.006, 0.035], P_f_ = 6.85×10^−3^). The allele with the largest estimated maternal and fetal effects was *B*39:06*, which had a relatively low allele frequency 0.69% and consequently wide confidence intervals for the effect sizes (maternal effect = -0.113 [-0.170, -0.057], P_m_ =8.49×10^−8^; fetal effect = 0.099 [0.004, 0.153], P_f_ = 4.03×10^−4^). We identified two nominally significant associations in the *HLA-C* gene, which has previously been reported to be associated with BW, including *HLA-C*03:03* (maternal effect = 0.038 [0.015, 0.061], P_m_ =1.26×10^−3^, fetal effect = -0.023 [-0.045, -0.001], P_f_ =4.21×10^−2^) and *HLA-C*04:01* (maternal effect = -0.027 [-0.047, -0.007], P_m_ =7.20×10^−3^). The most significant Class II allele was *DRB1*11:04*, which indicated only a fetal effect (fetal effect = −0.073 [−0.117, −0.029], P_f_ = 1.21×10^−3^). Supplementary Table S8 presents the results for all alleles and Supplementary Figure S5 presents the results from the SEM of the BW-associated HLA alleles at each locus.

**Figure 4.**
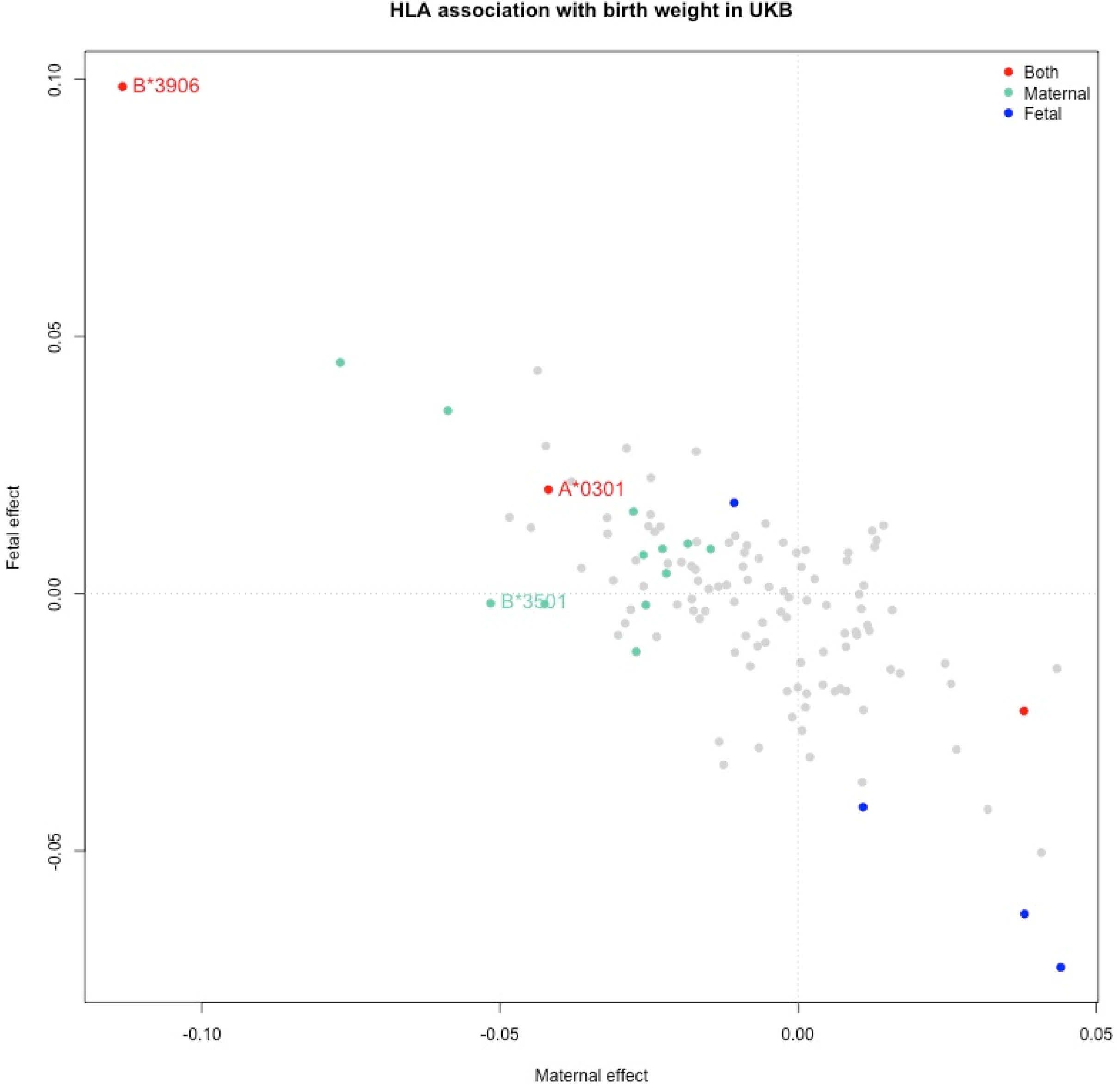
Maternal and fetal effect estimates for classical HLA alleles and BW in the UK Biobank Fetal and maternal effect sizes estimated using the structural equation model. The colour of each dot represents whether maternal (green) and/or fetal associations (blue) passed nominal significance (P < 0.05). Red dots indicate alleles where tests for both maternal and fetal effects reached the threshold. Light grey dots represent alleles which did not (P < 0.05). Alleles with P < 0.001 are explicitly labelled. The observed negative correlation between maternal and fetal effect estimates is at least partially a consequence of the negative correlation between these parameters in the SEM.

In conditional analyses including the previously reported BW associated SNPs in the MHC region (i.e. rs9366778 and rs6911024 ^8,12^) the effect of *HLA-B*35:01* (P_2df_ = 3.24×10^−3^, P_m_ = 1.83×10^−3^, P_f_ = 6.66×10^−2^) and *HLA-C*04:01* (P_2df_ = 0.221, P_m_ = 0.465, P_f_ = 0.626) were substantially attenuated in both two and one-degree of freedom tests (Supplementary Table S6 and S8). The rest of the top associated classical MHC alleles were still significant in two degrees of freedom test (P < 0.001), including *A*03:01, C*05:01, B*39:06*, and *DPB1*01:01*. The most significant allele was still *HLA-A*03:01* (P_m_ =1.17×10^−6^ and P_f_ = 1.71×10^−2^).

## Discussion

In the current study, we formulated a SEM for estimating the maternal and fetal effects of classical HLA alleles on perinatal phenotypes. Our SEM is appropriate for analyzing data structures where (unrelated) individuals with classical HLA genotypes are measured on their own phenotype and their offspring. Our simulations suggested that estimates of maternal and fetal allelic effects produced by our SEM were asymptotically unbiased. This was in contrast to analyses using ordinary least squares linear regression, which does not explicitly partition allelic effects into maternal and fetal components. Our SEM can also be used when a subset of the data has either only the individual’s own phenotype and/or only the phenotype of their offspring.

Asymptotic power calculations show that our SEM has appreciable power to partition maternal and fetal effects of common HLA alleles in a sample of at least 80,000 individuals (i.e. similar to the number of individuals in the UKB with BW information on themselves and their offspring). For example, we had greater than 80% power to partition the allele of interest into realistic maternal or fetal allelic effects (> 0.014 standardized effect size) assuming a sample size of 80,000 genotyped mothers who reported both their own and their offspring’s BW (α = 0.05, one degree of freedom test). As expected, the two degree of freedom test to detect any allelic effect at the locus (i.e. regardless of whether this effect was mediated by the maternal genome, the fetal genome, or both) had more power than the one degree of freedom test partitioning allelic effects into maternal or fetal components. The further addition of mother-offspring pairs with incomplete data (i.e. individuals who only report their own BW or mothers who only report their offspring’s BW) further increased the power of the one degree and two degree of freedom tests. These results are similar to what we have observed previously in our analogous SEM that analyses biallelic SNP genotypes^18,19^.

In empirical analyses in UKB, we identified BW associations at multiple alleles at MHC class I and II loci using data from more than 270,000 independent European individuals. We then used our SEM to partition the allelic associations into maternal and/or fetal effects. A total of 19 HLA alleles were shown to have maternal and/or fetal effects on BW (P < 0.05). Most classical HLA alleles which showed association with offspring BW, exhibited negative effects relative to the most common allele at the respective locus. Previous GWAS have shown that most genome-wide significant SNPs for birthweight in non-MHC regions have effects that are mediated primarily through the fetal genome (or at least the estimated fetal effect is larger than the estimated maternal effect) ^8,19^. However, in the HLA region, our results suggest that many alleles have predominantly maternal effects. In addition, our results suggest that the most common maternal alleles (i.e. which were used as the baseline comparator genotype) are protective against low birthweight and fetal growth restriction. It is interesting to speculate whether this could be a consequence of natural selection, since HLA alleles that cause low BW might lead to a low survival rate and consequently decrease their frequencies in the population.

The classical allele with the lowest p-value was *HLA-A*03:01*, which exhibited a predominantly maternal effect that decreased offspring BW relative to the most common *HLA-A*02:01* allele (P_2df_ =7.39 × 10^−8^, maternal effect = -0.042 [-0.057, -0.027], P_m_ =7.90×10^−8^)). The allele also exhibited a fetal effect (fetal effect = 0.0202 [0.0056, 0.0349], P_f_ = 6.85×10^−3^), but in the opposite direction to the maternal effect. The association was still significant after conditioning on genome-wide associated SNPs in the HLA region (P_2df_ =1.05 ×10^−6^).. HLA-*A*03:01* has never been directly associated with BW previously. However, it is associated with double the risk of developing multiple sclerosis (MS)^31^. Studies have shown that MS patients show a higher risk of high BW themselves than the general population^32^, while children who are born to mothers with MS might have lower BW for the same gestational age^33^. These directions are consistent with the direction of partitioned maternal and fetal effects of HLA-*A*03:01* on BW observed in our study. If *A*03:01* is truly associated with both phenotypes then this most likely reflects genetic pleiotropy rather than a causal effect mediated through MS. MS is a rare disease with a prevalence less than 0.5%^34^ and there are other classical HLA alleles with far stronger associations with MS, especially *DRB1*15:01*^35^, which would also be expected to show associations with BW if the effect were truly causal.

In addition to the putative association with the BW, *HLA-A* alleles are also in high linkage disequilibrium (LD) with non-classical HLA-Ib alleles (*HLA-E, -F* and *-G*), especially *HLA-G* ^36^. *HLA-G* plays a key role in immune tolerance throughout pregnancy, being primarily expressed by extravillous trophoblast cells in the placenta^37^. *HLA-G* genotypes have been associated with BW, placental weight ^38^, recurrent miscarriage ^39^ and pregnancy-induced hypertension^40^. It has been previously reported that the *HLA-A*03* allele is in strong linkage disequilibrium with *HLA-F*01:03:01*, the *HLA-G UTR-4* haplotype, and the *HLA-G*01:01* allele ^41^. The latter correlation has been further supported by a recent high-resolution study showing strong LD between *HLA-A*03:01:01:01* and *HLA-G*01:01:01:05*^42^. Persson et al. report that the maternal *UTR4-HLA-G*01:01:01:05* haplotype and the *F*01:03:01* allele were both associated with lower anti-HLA I complement activation and higher harmless IgG4 expression, which might reduce the risk of alloimmunization during pregnancy^43^. However, *HLA-A*03:01* shows a negative maternal effect on BW in our study, which is in contrast to the protective effect of *UTR4-HLA-G*01:01:01:05* haplotype and the HLA-*F*01:03:01* allele reported by the previous authors. However, their study only had a small sample size (N= 89) with genotyped maternal, but not fetal HLA alleles. Larger studies are required to elucidate the role of non-classical HLA Ib alleles with fetal growth restriction and alloimmunization during pregnancy.

The *HLA-B*39:06* allele displayed the largest estimated maternal and fetal effects among all associated alleles. A previous study has shown that the *HLA-B*39:06* allele is associated higher risk of type 1 diabetes (T1D)^44,45^, especially when two specific *HLA-DR/DQ* haplotypes are present (*DRB1*08:01-DQB1*04:02*, Odds Ratio (OR)=25.4; *DRB1*01:01-DQB1*05:01*, OR = 10.3)^46^. Intriguingly, *DRB1*08:01, DRB1*01:01* and *DQB1*05:01* were also nominally significantly associated with BW in our analysis (P_2df_ < 0.05). High BW has been previously associated with childhood-onset T1D ^47^ and pregnancies in women with T1D have also been associated with BW extremes and preterm delivery^48,49^. These associations could be partly attributable to *HLA-B*39:06* as our results indicate a positive fetal and a negative maternal effect of this allele on BW.

The most significant partitioned association in the class II region involved the *HLA-DRB1*11:04* allele which exhibited a significant fetal effect. Class II HLA molecules are not only present peptide antigens on the surface of antigen-precenting cells (APCs) for recognition by T helper cells, but they also play roles in the regulation of APCs and innate immunity. Our findings imply that class II HLA molecules may also be involved in intrauterine growth via the fetal immune response or feto-maternal immune interactions. Previous reports have shown an association between *HLA-DRB1*11* and early-onset diseases, including juvenile idiopathic arthritis^50^, and child-onset acquired thrombotic thrombocytopenic purpura^51^. It is possible that this allele may have important influences on growth and development in early life stages. The link between intrauterine exposures and early-onset diseases warrants further study.

The strength of the association between BW and both *HLA-C*04:01* and *-B*35:01* were significantly attenuated after conditioning on the BW associated SNPs rs9366778 and rs6911024^8^. This is to be expected given the linkage disequilibrium between the SNPs and these classical HLA alleles. For example, the SNP rs9366778 which exhibits a fetal effect^8^ on BW, is in low to moderate LD with *HLA-C*04:01* and *HLA-B*35:01* (r^2^ = 0.14, and 0.06, respectively; according to the reference panel from the Type 1 Diabetes Genetic Consortium reference panel ^52,53^). Likewise, the other SNP rs6911024 (*MICA*), which has a predominantly maternal effect^8^, is also in moderate LD with both *HLA-B*35:01* and *HLA-C*04:01* (r^2^ = 0.24 and 0.21, respectively, using the proxy SNP rs6910087 in the same reference panel [r^2^=0.99]^54^). The *HLA-B*35:01* and *HLA-C*04:01* alleles are also moderately correlated (r^2^ = 0.37). However, the *B*35:01* is still more significant (P_2df_ = 3.24×10^−3^) than either rs9366778 or rs6911024 in the conditional model (P_2df_ = 0.011 and 0.611, respectively) indicating the primary association at the classical *HLA-B* alleles rather than the SNPs. On the contrary, *C*0401* was totally conditioned by the two SNPs (P_2df_ = 0.221), while rs6911024 still shows nominal significance (P_2df_ = 2.81×10^−3^). The rest of the most associated alleles were still significant (*C*05:01,B*39:06*, and *DPB1*01:01*, P<0.001) indicating those independent associations might be novel findings of genetic contributions to fetal growth.

Fetal *HLA-C* epitopes have previously been associated with BW in observational studies^16,17^, and GWAS studies have reported a BW associated SNP rs9366778 in the *HLA-C* gene which exhibits a putative fetal effect^8^. We detected three HLA-C alleles in the two degrees of freedom test that reached nominal significance (*HLA-C*03:03, C*04:01, C*05:01*, P<0.05). In the one degree of freedom test, the only nominally significant fetal association at the HLA-C locus was found at *HLA-C*03:03* (fetal effect = – 0.228, P_f_= 0.042), but the allele shows stronger evidence for a maternal effect in the opposite direction (maternal effect = 0.379, P_m_= 0.0012). A previous study ^17^ has suggested the existence of an interaction between paternally derived fetal HLA-C2 epitope (HLA-C molecule has a lysine at position 80; *HLA-C*02/*04/*05/*06/*15*)^55^ and maternal killer-cell immunoglobulin-like receptor (KIR) genotypes influences human BW. There are only small numbers of genotyped-mother offspring pairs in the UKB and so we were unable to examine interaction effects between maternal and fetal genotypes in our study. Future studies involving genotyped parent-offspring trios and dyads are warranted.

There are several limitations to our study. First, our model is not capable of modelling maternal and fetal interactions or incompatibility in the HLA and KIR regions. Previous studies suggest that mother/offspring interactions between HLA and KIR genes may contribute to intrauterine growth restriction^16,17^. To investigate this hypothesis via genetic association would require large numbers of genotyped mother-offspring pairs/parent offspring trios, an alternative statistical model, which is an active area of research by our group. Second, we merged classical HLA alleles with low frequency (i.e. MAF < 0.5%) into a single category for computational efficiency and because we did not have enough power to resolve effects at low-frequency alleles. However, it is possible that rare alleles may affect BW and our pooling of alleles in this fashion may have obscured any such effects. Third, we evaluated HLA associations one locus at a time without modelling linkage disequilibrium or potential interactions between different HLA loci. Fourth, our findings do not exclude the possibility that the variants which functionally affect birthweight are in non-classical HLA genes or non-MHC genes and are in LD with the associated classical HLA alleles. Functional work may be the only way to resolve these different possibilities. Fifth, our analyses are based upon self-reported own and offspring BW in the UKB which may include varying degrees of measurement error. Nevertheless, in other cohorts, self-reported BW has shown to correlate well with hospital records ^56,57^. Sixth, gestational age was not reported in the UKB so could not be incorporated into our analyses. However, we excluded the lower and upper extremes of BW in our analyses to minimize potential influences from pre- or post-term births. Finally, we acknowledge that a major reason for our interest in classical HLA alleles and their possible relationship with BW is a consequence of genome-wide significant signals being found in the MHC region of the previous Early Growth Genetics consortium meta-analysis of birthweight^8^. As the UKB formed part of this sample, our results are not independent verification of the involvement of classical HLA alleles in BW etiology and consequently it will be important to replicate our findings in independent datasets in the future.

In conclusion, we have developed an SEM that can be used to partition the genetic association between classical HLA alleles and perinatal traits into maternal and fetal genetic components. Application of our model to individuals in the UKB revealed interesting novel allelic associations between BW and classical HLA alleles which potentially provide insight into the immunogenetics of intrauterine fetal growth.

## Acknowledgments

This research has been conducted using the UK Biobank resource (Reference 53641). G.W. is supported by the University of Queensland Graduate School Scholarship (Australian Government Research Training Program Scholarship). N.M.W. is funded by a National Health and Medical Research Council (Australia) Investigator grant (APP2008723). D.M.E. is funded by an Australian National Health and Medical Research Council Senior Research Fellowship (APP1137714) and this work was funded by NHMRC project grants (GNT1157714, GNT1183074). The authors would like to thank Dr John P. Kemp (Institute for Molecular Biosciences, The University of Queensland, Brisbane, Australia) for his kind assistance with the definition of European ancestry.

## Author contributions

G.W. conducted the statistical analysis and drafted the manuscript. G.W., N.M.W., and D.M.E. designed the study, interpreted the results and reviewed the manuscript.

## Competing interests

None

## Materials & Correspondence

Correspondence and requests for materials should be addressed to G.W. or D.M.E.

## Data availability

Human genotype and phenotype data from the UKB on which the results of this study were based were accessed with accession ID 53641. The genotype and phenotype data are available upon application to the UKB (http://www.ukbiobank.ac.uk/).

## Notes

### Competing Interest Statement

The authors have declared no competing interest.

### Summary of Updates

Data availability and acknowledgments updated.

