## Supplementary Materials for "Partitioning genetic effects on birth weight at classical human leukocyte antigen loci into indirect maternal and direct fetal components using structural equation modelling"

Supplementary Table S1-8

Table S1: Bias in estimating the effect of the HLA-X1 allele on birth weight in the simulation study, using both the structural equation model (SEM) and a linear model with varying maternal and fetal effect sizes, where all 80,000 simulated individuals reported both their own and their offspring's birth weight and the allele frequency HLA-X1 was 10%.

| **Simulated fetal effects** | | | **Simulated maternal effects** | | | **Bias** | | | |
|---|---|---|---|---|---|---|---|---|---|
| **Allele 1 ($\beta_{f1}$)** | **Allele 2 ($\beta_{f2}$)** | **Allele 3 ($\beta_{f3}$)** | **Allele 1 ($\beta m_1$)** | **Allele 2 ($\beta m_2$)** | **Allele 3 ($\beta m_3$)** | **Linear model - fetal effect (95% CI)** | **Linear model - maternal effect (95% CI)** | **Structural equation model - fetal effect (95% CI)** | **Structural equation model - maternal effect (95% CI)** |
| 0 | 0.014 | 0.02 | -0.01 | 0 | -0.014 | -8.02E-03 ( -8.03E-03 , -8.02E-03) | -3.54E-04 ( -3.61E-04 , -3.47E-04) | 3.38E-06 ( -5.75E-06 , 1.25E-05) | -1.92E-05 ( -2.88E-05 , -9.61E-06) |
| 0 | 0.014 | 0.02 | -0.014 | 0 | -0.01 | -1.02E-02 ( -1.03E-02 , -1.02E-02) | -7.99E-04 ( -8.06E-04 , -7.92E-04) | 3.42E-06 ( -5.71E-06 , 1.25E-05) | -1.92E-05 ( -2.88E-05 , -9.64E-06) |
| 0 | 0.014 | 0.02 | 0 | -0.01 | -0.014 | -2.47E-03 ( -2.47E-03 , -2.46E-03) | 7.56E-04 ( 7.49E-04 , 7.63E-04) | 4.60E-06 ( -4.54E-06 , 1.37E-05) | -1.98E-05 ( -2.94E-05 , -1.03E-05) |
| 0 | 0.014 | 0.02 | 0 | 0.01 | 0.014 | -5.14E-03 ( -5.14E-03 , -5.13E-03) | -4.58E-03 ( -4.58E-03 , -4.57E-03) | -4.46E-07 ( -9.58E-06 , 8.68E-06) | -1.73E-05 ( -2.68E-05 , -7.70E-06) |
| 0 | 0.014 | 0.02 | 0.01 | 0 | 0.014 | 4.20E-04 ( 4.14E-04 , 4.27E-04) | -3.47E-03 ( -3.47E-03 , -3.46E-03) | 8.05E-07 ( -8.33E-06 , 9.94E-06) | -1.79E-05 ( -2.75E-05 , -8.35E-06) |
| 0 | 0.014 | 0.02 | 0.014 | 0 | 0.01 | 2.64E-03 ( 2.64E-03 , 2.65E-03) | -3.02E-03 ( -3.03E-03 , -3.02E-03) | 7.48E-07 ( -8.39E-06 , 9.89E-06) | -1.79E-05 ( -2.75E-05 , -8.30E-06) |
| 0.014 | 0 | 0.02 | -0.01 | 0 | -0.014 | -6.47E-03 ( -6.47E-03 , -6.46E-03) | 7.42E-03 ( 7.42E-03 , 7.43E-03) | 3.87E-06 ( -5.27E-06 , 1.30E-05) | -2.01E-05 ( -2.97E-05 , -1.06E-05) |
| 0.014 | 0 | 0.02 | -0.014 | 0 | -0.01 | -8.69E-03 ( -8.70E-03 , -8.68E-03) | 6.98E-03 ( 6.97E-03 , 6.99E-03) | 3.85E-06 ( -5.28E-06 , 1.30E-05) | -2.01E-05 ( -2.97E-05 , -1.05E-05) |
| 0.014 | 0 | 0.02 | 0 | -0.01 | -0.014 | -9.11E-04 ( -9.18E-04 , -9.05E-04) | 8.53E-03 ( 8.53E-03 , 8.54E-03) | 5.02E-06 ( -4.12E-06 , 1.42E-05) | -2.07E-05 ( -3.03E-05 , -1.11E-05) |
| 0.014 | 0 | 0.02 | 0 | 0.01 | 0.014 | -3.58E-03 ( -3.59E-03 , -3.57E-03) | 3.20E-03 ( 3.19E-03 , 3.21E-03) | 5.04E-08 ( -9.08E-06 , 9.18E-06) | -1.82E-05 ( -2.78E-05 , -8.65E-06) |
| 0.014 | 0 | 0.02 | 0.01 | 0 | 0.014 | 1.98E-03 ( 1.97E-03 , 1.98E-03) | 4.31E-03 ( 4.30E-03 , 4.32E-03) | 1.24E-06 ( -7.90E-06 , 1.04E-05) | -1.88E-05 ( -2.84E-05 , -9.25E-06) |
| 0.014 | 0 | 0.02 | 0.014 | 0 | 0.01 | 4.20E-03 ( 4.19E-03 , 4.20E-03) | 4.75E-03 ( 4.75E-03 , 4.76E-03) | 1.28E-06 ( -7.86E-06 , 1.04E-05) | -1.88E-05 ( -2.84E-05 , -9.27E-06) |
| 0.02 | 0.014 | 0 | -0.01 | 0 | -0.014 | -5.80E-03 ( -5.81E-03 , -5.79E-03) | 1.08E-02 ( 1.08E-02 , 1.08E-02) | 3.05E-06 ( -6.09E-06 , 1.22E-05) | -1.85E-05 ( -2.80E-05 , -8.89E-06) |

| | | | | | | | | | |
|---|---|---|---|---|---|---|---|---|---|
| 0.02 | 0.014 | 0 | -0.014 | 0 | -0.01 | -8.02E-03 ( -8.03E-03 , -8.01E-03) | 1.03E-02 ( 1.03E-02 , 1.03E-02) | 3.03E-06 ( -6.11E-06 , 1.22E-05) | -1.84E-05 ( -2.80E-05 , -8.87E-06) |
| 0.02 | 0.014 | 0 | 0 | -0.01 | -0.014 | -2.42E-04 ( -2.49E-04 , -2.36E-04) | 1.19E-02 ( 1.19E-02 , 1.19E-02) | 4.17E-06 ( -4.96E-06 , 1.33E-05) | -1.90E-05 ( -2.86E-05 , -9.44E-06) |
| 0.02 | 0.014 | 0 | 0 | 0.01 | 0.014 | -2.91E-03 ( -2.92E-03 , -2.91E-03) | 6.53E-03 ( 6.53E-03 , 6.54E-03) | -7.78E-07 ( -9.92E-06 , 8.36E-06) | -1.65E-05 ( -2.61E-05 , -6.96E-06) |
| 0.02 | 0.014 | 0 | 0.01 | 0 | 0.014 | 2.64E-03 ( 2.64E-03 , 2.65E-03) | 7.65E-03 ( 7.64E-03 , 7.65E-03) | 4.07E-07 ( -8.73E-06 , 9.55E-06) | -1.71E-05 ( -2.67E-05 , -7.57E-06) |
| 0.02 | 0.014 | 0 | 0.014 | 0 | 0.01 | 4.87E-03 ( 4.86E-03 , 4.87E-03) | 8.09E-03 ( 8.08E-03 , 8.10E-03) | 4.10E-07 ( -8.73E-06 , 9.55E-06) | -1.71E-05 ( -2.67E-05 , -7.56E-06) |

βm : maternal effect; βf : fetal effect; The “1”, “2”, and “3” subscripts correspond to the classical HLA alleles HLA-X1, HLA-X2, and HLA-X3. The “m” and “f” subscripts refer to estimates of the maternal and fetal effect, respectively. The slope of the linear regression of offspring birthweight on own genotype is used as an estimate for the maternal effect, and the slope of the linear regression of own birthweight on own genotype is used as an estimate of the fetal effect. The allele frequencies of all alternative alleles (X1, X2, and X3) were set to 10%.

Table S2: Type 1 error rate and power to detect a significant effect of the HLA-X1 allele ($\alpha = 0.05$) in the simulation study, using both the structural equation model (SEM) and a linear model with varying maternal and fetal effect sizes, where all 80,000 simulated individuals reported both their own and their offspring's birth weight and the allele frequency HLA-X1 was 10%.

| **Allele 1 ($\beta f_1$)** | **Allele 2 ($\beta f_2$)** | **Allele 3 ($\beta f_3$)** | **Allele 1 ($\beta m_1$)** | **Allele 2 ($\beta m_2$)** | **Allele 3 ($\beta m_3$)** | **Linear model - fetal effect (95% CI)** | **Structural equation model - fetal effect (95% CI)** | **Linear model - maternal effect (95% CI)** | **Structural equation model - maternal effect (95% CI)** |
|---|---|---|---|---|---|---|---|---|---|
| 0 | 0.014 | 0.02 | -0.01 | 0 | -0.014 | 0.626 (0.596, 0.656) | 0.044 (0.031, 0.057) | 0.825 (0.801, 0.849) | 0.547 (0.516, 0.578) |
| 0 | 0.014 | 0.02 | -0.014 | 0 | -0.01 | 0.836 (0.813, 0.859) | 0.045 (0.032, 0.058) | 0.987 (0.98, 0.994) | 0.828 (0.805, 0.851) |
| 0 | 0.014 | 0.02 | 0 | -0.01 | -0.014 | 0.09 (0.072, 0.108) | 0.043 (0.03, 0.056) | 0.064 (0.049, 0.079) | 0.053 (0.039, 0.067) |
| 0 | 0.014 | 0.02 | 0 | 0.01 | 0.014 | 0.303 (0.275, 0.331) | 0.045 (0.032, 0.058) | 0.251 (0.224, 0.278) | 0.052 (0.038, 0.066) |

| | | | | | | | | | |
|---|---|---|---|---|---|---|---|---|---|
| 0 | 0.014 | 0.02 | 0.01 | 0 | 0.014 | 0.04 (0.028, 0.052) | 0.044 (0.031, 0.057) | 0.453 (0.422, 0.484) | 0.549 (0.518, 0.58) |
| 0 | 0.014 | 0.02 | 0.014 | 0 | 0.01 | 0.098 (0.08, 0.116) | 0.044 (0.031, 0.057) | 0.882 (0.862, 0.902) | 0.819 (0.795, 0.843) |
| 0.014 | 0 | 0.02 | -0.01 | 0 | -0.014 | 0.57 (0.539, 0.601) | 0.855 (0.833, 0.877) | 0.106 (0.087, 0.125) | 0.547 (0.516, 0.578) |
| 0.014 | 0 | 0.02 | -0.014 | 0 | -0.01 | 0.31 (0.281, 0.339) | 0.855 (0.833, 0.877) | 0.515 (0.484, 0.546) | 0.827 (0.804, 0.85) |
| 0.014 | 0 | 0.02 | 0 | -0.01 | -0.014 | 0.973 (0.963, 0.983) | 0.851 (0.829, 0.873) | 0.664 (0.635, 0.693) | 0.053 (0.039, 0.067) |
| 0.014 | 0 | 0.02 | 0 | 0.01 | 0.014 | 0.839 (0.816, 0.862) | 0.847 (0.825, 0.869) | 0.155 (0.133, 0.177) | 0.052 (0.038, 0.066) |
| 0.014 | 0 | 0.02 | 0.01 | 0 | 0.014 | 0.997 (0.994, 1) | 0.846 (0.824, 0.868) | 0.978 (0.969, 0.987) | 0.549 (0.518, 0.58) |
| 0.014 | 0 | 0.02 | 0.014 | 0 | 0.01 | 1 (1, 1) | 0.848 (0.826, 0.87) | 0.999 (0.997, 1.001) | 0.822 (0.798, 0.846) |
| 0.02 | 0.014 | 0 | -0.01 | 0 | -0.014 | 0.987 (0.98, 0.994) | 0.989 (0.983, 0.995) | 0.066 (0.051, 0.081) | 0.55 (0.519, 0.581) |
| 0.02 | 0.014 | 0 | -0.014 | 0 | -0.01 | 0.944 (0.93, 0.958) | 0.989 (0.983, 0.995) | 0.177 (0.153, 0.201) | 0.83 (0.807, 0.853) |
| 0.02 | 0.014 | 0 | 0 | -0.01 | -0.014 | 1 (1, 1) | 0.984 (0.976, 0.992) | 0.921 (0.904, 0.938) | 0.052 (0.038, 0.066) |
| 0.02 | 0.014 | 0 | 0 | 0.01 | 0.014 | 1 (1, 1) | 0.978 (0.969, 0.987) | 0.445 (0.414, 0.476) | 0.051 (0.037, 0.065) |
| 0.02 | 0.014 | 0 | 0.01 | 0 | 0.014 | 1 (1, 1) | 0.988 (0.981, 0.995) | 0.999 (0.997, 1.001) | 0.55 (0.519, 0.581) |
| 0.02 | 0.014 | 0 | 0.014 | 0 | 0.01 | 1 (1, 1) | 0.988 (0.981, 0.995) | 1 (1, 1) | 0.825 (0.801, 0.849) |

βm : maternal effect; βf : fetal effect; The “1”, “2”, and “3” subscripts correspond to the classical HLA alleles HLA-X1, HLA-X2, and HLA-X3. The “m” and “f” subscripts refer to estimates of the maternal and fetal effect, respectively. The slope of the linear regression of offspring birthweight on own genotype is used as an estimate for the maternal effect, and the slope of the linear regression of own birthweight on own genotype is used as an estimate of the fetal effect. The allele frequencies of all alternative alleles (X1, X2, and X3) were set to 10%. The pink highlight indicates the corresponding type 1 errors in SEM when the true effect of the allele of interest were set to zero.

Table S3: Asymptotic power to detect maternal (βm1) and fetal (βf1) effects of HLA-X1 allele on birth weight (α = 0.05), where all 80,000 simulated individuals reported both their own and their offspring's birth weight and the allele frequency HLA-X1 was 10%.

| Fetal effects | | | Maternal effects | | | Power | | |
|---|---|---|---|---|---|---|---|---|
| **Allele 1 ($\beta_{f1}$)** | **Allele 2 ($\beta_{f2}$)** | **Allele 3 ($\beta_{f3}$)** | **Allele 1 ($\beta m_1$)** | **Allele 2 ($\beta m_2$)** | **Allele 3 ($\beta m_3$)** | **Fetal** | **Maternal** | **Both** |
| 0 | 0.014 | 0.02 | -0.01 | 0 | -0.014 | 5.00% | 55.25% | 82.14% |
| 0 | 0.014 | 0.02 | -0.014 | 0 | -0.01 | 5.00% | 83.37% | 98.09% |
| 0 | 0.014 | 0.02 | 0 | -0.01 | -0.014 | 5.00% | 5.00% | 5.00% |
| 0 | 0.014 | 0.02 | 0 | 0.01 | 0.014 | 5.00% | 5.00% | 5.00% |
| 0 | 0.014 | 0.02 | 0.01 | 0 | 0.014 | 5.00% | 55.28% | 82.15% |
| 0 | 0.014 | 0.02 | 0.014 | 0 | 0.01 | 5.00% | 83.38% | 98.09% |
| 0.014 | 0 | 0.02 | -0.01 | 0 | -0.014 | 83.36% | 55.24% | 83.41% |
| 0.014 | 0 | 0.02 | -0.014 | 0 | -0.01 | 83.36% | 83.36% | 89.03% |
| 0.014 | 0 | 0.02 | 0 | -0.01 | -0.014 | 83.37% | 5.00% | 98.09% |
| 0.014 | 0 | 0.02 | 0 | 0.01 | 0.014 | 83.39% | 5.00% | 98.09% |
| 0.014 | 0 | 0.02 | 0.01 | 0 | 0.014 | 83.40% | 55.29% | 100.00% |
| 0.014 | 0 | 0.02 | 0.014 | 0 | 0.01 | 83.40% | 83.39% | 100.00% |
| 0.02 | 0.014 | 0 | -0.01 | 0 | -0.014 | 98.69% | 55.25% | 99.09% |
| 0.02 | 0.014 | 0 | -0.014 | 0 | -0.01 | 98.69% | 83.36% | 98.69% |
| 0.02 | 0.014 | 0 | 0 | -0.01 | -0.014 | 98.69% | 5.00% | 99.99% |
| 0.02 | 0.014 | 0 | 0 | 0.01 | 0.014 | 98.70% | 5.00% | 99.99% |
| 0.02 | 0.014 | 0 | 0.01 | 0 | 0.014 | 98.70% | 55.27% | 100.00% |
| 0.02 | 0.014 | 0 | 0.014 | 0 | 0.01 | 98.70% | 83.39% | 100.00% |

βm : maternal effect; βf : fetal effect; The "1", "2", and "3" subscripts correspond to the classical HLA alleles HLA-X1, HLA-X2, and HLA-X3. "Maternal", "Fetal", and "Both" refer to testing maternal, fetal, and both effects. The allele frequencies of all alternative alleles (X1, X2, and X3) were set to 10%.

Table S4: Asymptotic power to detect maternal (βm1) and fetal (βf1) effects of HLA-X1 allele on birth weight ($\alpha = 0.05$), where 80,000 individuals have both own and offpsring birth weight, 100,000 individual have their own birth weight only, and 80,000 individuals have birth weight of their offspring only and the allele frequency HLA-X1 was 10%.

| **Fetal effects** | | | **Maternal effects** | | | **Power** | | |
|---|---|---|---|---|---|---|---|---|
| **Allele 1 ($\beta_{f1}$)** | **Allele 2 ($\beta_{f2}$)** | **Allele 3 ($\beta_{f3}$)** | **Allele 1 ($\beta m_1$)** | **Allele 2 ($\beta m_2$)** | **Allele 3 ($\beta m_3$)** | **Fetal** | **Maternal** | **Both** |
| 0 | 0.014 | 0.02 | -0.01 | 0 | -0.014 | 5.00% | 80.94% | 99.04% |
| 0 | 0.014 | 0.02 | -0.014 | 0 | -0.01 | 5.00% | 97.78% | 100.00% |
| 0 | 0.014 | 0.02 | 0 | -0.01 | -0.014 | 5.00% | 5.00% | 5.00% |
| 0 | 0.014 | 0.02 | 0 | 0.01 | 0.014 | 5.00% | 5.00% | 5.00% |
| 0 | 0.014 | 0.02 | 0.01 | 0 | 0.014 | 5.00% | 80.96% | 99.04% |
| 0 | 0.014 | 0.02 | 0.014 | 0 | 0.01 | 5.00% | 97.78% | 100.00% |
| 0.014 | 0 | 0.02 | -0.01 | 0 | -0.014 | 98.47% | 80.93% | 98.55% |
| 0.014 | 0 | 0.02 | -0.014 | 0 | -0.01 | 98.47% | 97.78% | 99.11% |
| 0.014 | 0 | 0.02 | 0 | -0.01 | -0.014 | 98.48% | 5.00% | 100.00% |
| 0.014 | 0 | 0.02 | 0 | 0.01 | 0.014 | 98.48% | 5.00% | 100.00% |
| 0.014 | 0 | 0.02 | 0.01 | 0 | 0.014 | 98.48% | 80.97% | 100.00% |

| | | | | | | | | |
|---|---|---|---|---|---|---|---|---|
| 0.014 | 0 | 0.02 | 0.014 | 0 | 0.01 | 98.48% | 97.79% | 100.00% |
| 0.02 | 0.014 | 0 | -0.01 | 0 | -0.014 | 100.00% | 80.95% | 100.00% |
| 0.02 | 0.014 | 0 | -0.014 | 0 | -0.01 | 100.00% | 97.78% | 100.00% |
| 0.02 | 0.014 | 0 | 0 | -0.01 | -0.014 | 100.00% | 5.00% | 100.00% |
| 0.02 | 0.014 | 0 | 0 | 0.01 | 0.014 | 100.00% | 5.00% | 100.00% |
| 0.02 | 0.014 | 0 | 0.01 | 0 | 0.014 | 100.00% | 80.96% | 100.00% |
| 0.02 | 0.014 | 0 | 0.014 | 0 | 0.01 | 100.00% | 97.78% | 100.00% |

βm : maternal effect; βf : fetal effect; The “1”, “2”, and “3” subscripts correspond to the classical HLA alleles HLA-X1, HLA-X2, and HLA-X3. “Maternal”, “Fetal”, and “Both” refer to testing maternal, fetal, and both effects. The allele frequencies of all alternative alleles (X1, X2, and X3) were set to 10%.

Table S5: Details of the frequency of imputed alleles at each classical HLA loci in the UK Biobank and how they were included in the analysis

| **Locus** | **Allele** | **Frequency** | **Allele Included in Analysis** |
|---|---|---|---|
| A | A*0101 | 19.4453% | Included |
| | A*0102 | 0.0209% | Others |
| | A*0103 | 0.0024% | Others |
| | A*0201 | 27.6270% | Baseline |
| | A*0202 | 0.0679% | Others |
| | A*0203 | 0.1604% | Others |
| | A*0205 | 0.7506% | Included |
| | A*0206 | 0.1782% | Others |
| | A*0207 | 0.0035% | Others |

| | | |
|---|---|---|
| A*0211 | 0.0985% | Others |
| A*0220 | 0.0075% | Others |
| A*0301 | 14.5062% | Included |
| A*0302 | 0.1182% | Others |
| A*1101 | 6.0632% | Included |
| A*1102 | 0.0026% | Others |
| A*2301 | 1.6939% | Included |
| A*2402 | 7.3721% | Included |
| A*2403 | 0.0513% | Others |
| A*2404 | 0.0029% | Others |
| A*2407 | 0.0077% | Others |
| A*2501 | 1.7308% | Included |
| A*2601 | 1.9735% | Included |
| A*2608 | 0.0717% | Others |
| A*2901 | 0.1253% | Others |
| A*2902 | 4.2392% | Included |
| A*3001 | 0.9959% | Included |
| A*3002 | 0.9765% | Included |
| A*3004 | 0.0942% | Others |
| A*3101 | 2.6519% | Included |
| A*3201 | 3.4632% | Included |
| A*3301 | 0.5402% | Included |
| A*3303 | 0.1815% | Others |
| A*3402 | 0.0884% | Others |
| A*3601 | 0.0011% | Others |
| A*6601 | 0.2847% | Others |
| A*6801 | 3.0023% | Included |
| A*6802 | 0.5911% | Included |

| | | | |
|---|---|---|---|
| | A*6817 | 0.0015% | Others |
| | A*6901 | 0.0287% | Others |
| | A*7401 | 0.0366% | Others |
| | A*7403 | 0.0040% | Others |
| | A*8001 | 0.0062% | Others |
| B | B*0702 | 14.9442% | Baseline |
| | B*0705 | 0.1350% | Others |
| | B*0801 | 14.4483% | Included |
| | B*1301 | 0.0013% | Others |
| | B*1302 | 1.9199% | Included |
| | B*1401 | 1.1913% | Included |
| | B*1402 | 2.5481% | Included |
| | B*1501 | 6.2069% | Included |
| | B*1502 | 0.0022% | Others |
| | B*1503 | 0.0660% | Others |
| | B*1505 | 0.0011% | Others |
| | B*1507 | 0.0080% | Others |
| | B*1509 | 0.0011% | Others |
| | B*1510 | 0.0143% | Others |
| | B*1516 | 0.0362% | Others |
| | B*1517 | 0.1709% | Others |
| | B*1518 | 0.3093% | Others |
| | B*1520 | 0.0015% | Others |
| | B*1524 | 0.0293% | Others |
| | B*1525 | 0.0020% | Others |
| | B*1801 | 3.6723% | Included |
| | B*1803 | 0.0192% | Others |
| | B*2702 | 0.1375% | Others |

| | | |
|---|---|---|
| B*2703 | 0.0070% | Others |
| B*2705 | 3.9501% | Included |
| B*2707 | 0.0071% | Others |
| B*3501 | 4.6762% | Included |
| B*3502 | 0.2055% | Others |
| B*3503 | 0.8526% | Included |
| B*3505 | 0.0053% | Others |
| B*3508 | 0.1354% | Others |
| B*3512 | 0.0055% | Others |
| B*3522 | 0.0011% | Others |
| B*3701 | 1.3772% | Included |
| B*3801 | 0.7585% | Included |
| B*3901 | 0.7762% | Included |
| B*3902 | 0.0018% | Others |
| B*3906 | 0.6859% | Included |
| B*3910 | 0.0060% | Others |
| B*4001 | 5.5216% | Included |
| B*4002 | 0.8323% | Included |
| B*4006 | 0.0287% | Others |
| B*4009 | 0.0009% | Others |
| B*4012 | 0.0013% | Others |
| B*4101 | 0.1956% | Others |
| B*4102 | 0.3450% | Others |
| B*4201 | 0.0152% | Others |
| B*4402 | 11.2159% | Included |
| B*4403 | 5.8891% | Included |
| B*4404 | 0.0538% | Others |
| B*4405 | 0.1298% | Others |

| | | |
|---|---|---|
| B*4501 | 0.6457% | Included |
| B*4601 | 0.0022% | Others |
| B*4701 | 0.2982% | Others |
| B*4801 | 0.0143% | Others |
| B*4901 | 1.1426% | Included |
| B*5001 | 0.8460% | Included |
| B*5002 | 0.0011% | Others |
| B*5101 | 3.6586% | Included |
| B*5102 | 0.0011% | Others |
| B*5105 | 0.0024% | Others |
| B*5106 | 0.0011% | Others |
| B*5107 | 0.0104% | Others |
| B*5108 | 0.0721% | Others |
| B*5201 | 0.5106% | Included |
| B*5301 | 0.2406% | Others |
| B*5401 | 0.0029% | Others |
| B*5501 | 1.8252% | Included |
| B*5502 | 0.0020% | Others |
| B*5601 | 0.3491% | Others |
| B*5604 | 0.0013% | Others |
| B*5701 | 3.9154% | Included |
| B*5702 | 0.0216% | Others |
| B*5703 | 0.0070% | Others |
| B*5801 | 0.4935% | Others |
| B*5802 | 0.0026% | Others |
| B*7301 | 0.0159% | Others |
| B*7801 | 0.0013% | Others |
| B*8101 | 0.0016% | Others |

| | | | |
|---|---|---|---|
| C | C*0102 | 3.3953% | Included |
| | C*0202 | 3.5985% | Included |
| | C*0302 | 0.1301% | Others |
| | C*0303 | 5.5702% | Included |
| | C*0304 | 7.9600% | Included |
| | C*0401 | 8.3180% | Included |
| | C*0403 | 0.0051% | Others |
| | C*0501 | 11.3025% | Included |
| | C*0602 | 8.9732% | Included |
| | C*0701 | 17.6493% | Others |
| | C*0702 | 15.9275% | Included |
| | C*0704 | 1.7523% | Included |
| | C*0801 | 0.0152% | Others |
| | C*0802 | 3.6693% | Included |
| | C*1202 | 0.4954% | Others |
| | C*1203 | 2.8865% | Included |
| | C*1402 | 0.9109% | Included |
| | C*1502 | 1.7396% | Included |
| | C*1504 | 0.0143% | Others |
| | C*1505 | 0.1223% | Others |
| | C*1601 | 4.4570% | Included |
| | C*1602 | 0.1721% | Others |
| | C*1604 | 0.0573% | Others |
| | C*1701 | 0.5325% | Included |
| | C*1801 | 0.0097% | Others |
| DRB1 | DRB1*0101 | 9.3538% | Included |
| | DRB1*0102 | 0.7226% | Included |
| | DRB1*0103 | 1.7395% | Included |

| | | |
|---|---|---|
| DRB1*0301 | 14.8943% | Baseline |
| DRB1*0302 | 0.0035% | Others |
| DRB1*0401 | 11.4300% | Included |
| DRB1*0402 | 0.2872% | Others |
| DRB1*0403 | 0.2177% | Others |
| DRB1*0404 | 4.0473% | Included |
| DRB1*0405 | 0.4772% | Others |
| DRB1*0406 | 0.0362% | Others |
| DRB1*0407 | 0.9805% | Included |
| DRB1*0408 | 0.0708% | Others |
| DRB1*0410 | 0.0024% | Others |
| DRB1*0411 | 0.0020% | Others |
| DRB1*0701 | 14.6397% | Included |
| DRB1*0801 | 1.8165% | Included |
| DRB1*0802 | 0.0315% | Others |
| DRB1*0803 | 0.2442% | Others |
| DRB1*0804 | 0.0827% | Others |
| DRB1*0806 | 0.0520% | Others |
| DRB1*0901 | 1.2817% | Included |
| DRB1*1001 | 0.5924% | Included |
| DRB1*1101 | 3.4985% | Included |
| DRB1*1102 | 0.1861% | Others |
| DRB1*1103 | 0.3101% | Others |
| DRB1*1104 | 1.1619% | Included |
| DRB1*1111 | 0.0013% | Others |
| DRB1*1201 | 1.4484% | Included |
| DRB1*1202 | 0.0049% | Others |
| DRB1*1301 | 5.0520% | Included |

| | | | |
|---|---|---|---|
| | DRB1*1302 | 3.8194% | Included |
| | DRB1*1303 | 0.8437% | Included |
| | DRB1*1304 | 0.0066% | Others |
| | DRB1*1305 | 0.0077% | Others |
| | DRB1*1312 | 0.0031% | Others |
| | DRB1*1321 | 0.0011% | Others |
| | DRB1*1401 | 1.9175% | Included |
| | DRB1*1402 | 0.0095% | Others |
| | DRB1*1403 | 0.0009% | Others |
| | DRB1*1404 | 0.0634% | Others |
| | DRB1*1406 | 0.0024% | Others |
| | DRB1*1409 | 0.0009% | Others |
| | DRB1*1415 | 0.0024% | Others |
| | DRB1*1501 | 14.6156% | Included |
| | DRB1*1502 | 0.4162% | Others |
| | DRB1*1503 | 0.0168% | Others |
| | DRB1*1601 | 0.5535% | Included |
| | DRB1*1602 | 0.0375% | Others |
| DRB3 | DRB3*0101 | 16.7939% | Included |
| | DRB3*0202 | 13.3762% | Included |
| | DRB3*0210 | 0.0026% | Not included |
| | DRB3*0224 | 0.0093% | Not included |
| | DRB3*0301 | 3.8177% | Included |
| | DRB3*9901 | 65.8670% | Baseline |
| DRB4 | DRB4*0101 | 8.3402% | Included |
| | DRB4*0103 | 25.9413% | Included |
| | DRB4*9901 | 65.3159% | Baseline |
| DRB5 | DRB5*0101 | 14.5655% | Included |

| | | | |
|---|---|---|---|
| | DRB5*0202 | 0.7135% | Included |
| | DRB5*9901 | 84.7110% | Baseline |
| DQB1 | DQB1*0201 | 15.0023% | Included |
| | DQB1*0202 | 10.4474% | Included |
| | DQB1*0301 | 17.5524% | Baseline |
| | DQB1*0302 | 10.2673% | Included |
| | DQB1*0303 | 5.2303% | Included |
| | DQB1*0304 | 0.1198% | Others |
| | DQB1*0401 | 0.0055% | Others |
| | DQB1*0402 | 1.9892% | Included |
| | DQB1*0501 | 12.0130% | Included |
| | DQB1*0502 | 0.6934% | Included |
| | DQB1*0503 | 2.1513% | Included |
| | DQB1*0504 | 0.1454% | Others |
| | DQB1*0601 | 0.4226% | Others |
| | DQB1*0602 | 14.4185% | Included |
| | DQB1*0603 | 5.1615% | Included |
| | DQB1*0604 | 2.7539% | Included |
| | DQB1*0605 | 0.0049% | Others |
| | DQB1*0609 | 1.0431% | Included |
| DQA1 | DQA1*0101 | 14.1357% | Included |
| | DQA1*0102 | 19.3068% | Included |
| | DQA1*0103 | 5.4948% | Included |
| | DQA1*0104 | 0.0048% | Others |
| | DQA1*0201 | 14.6568% | Included |
| | DQA1*0301 | 20.2463% | Included |
| | DQA1*0302 | 0.0053% | Others |
| | DQA1*0303 | 0.1144% | Others |

| | | | |
|---|---|---|---|
| | DQA1*0401 | 1.9410% | Included |
| | DQA1*0501 | 23.2705% | Baseline |
| | DQA1*0503 | 0.0035% | Others |
| | DQA1*0505 | 0.1128% | Others |
| | DQA1*0509 | 0.0062% | Others |
| | DQA1*0601 | 0.2591% | Others |
| DPB1 | DPB1*0101 | 5.9102% | Included |
| | DPB1*0201 | 11.1824% | Included |
| | DPB1*0202 | 0.7575% | Included |
| | DPB1*0301 | 10.3313% | Included |
| | DPB1*0401 | 44.5516% | Baseline |
| | DPB1*0402 | 10.7681% | Included |
| | DPB1*0501 | 2.0901% | Included |
| | DPB1*0601 | 1.2131% | Included |
| | DPB1*0901 | 0.5456% | Included |
| | DPB1*1001 | 1.6585% | Included |
| | DPB1*1101 | 2.6264% | Included |
| | DPB1*1301 | 1.4885% | Included |
| | DPB1*1401 | 0.9237% | Included |
| | DPB1*1501 | 0.7411% | Included |
| | DPB1*1601 | 0.4849% | Others |
| | DPB1*1701 | 1.1192% | Included |
| | DPB1*1801 | 0.0031% | Others |
| | DPB1*1901 | 0.5234% | Included |
| | DPB1*2001 | 0.0988% | Others |
| | DPB1*2101 | 0.0020% | Others |
| | DPB1*2301 | 0.1235% | Others |
| | DPB1*2601 | 0.0146% | Others |

| | | | |
|---|---|---|---|
| | DPB1*3401 | 0.0018% | Others |
| DPA1 | DPA1*0103 | 81.6168% | Baseline |
| | DPA1*0104 | 0.5189% | Included |
| | DPA1*0105 | 0.0562% | Not included |
| | DPA1*0201 | 14.6015% | Included |
| | DPA1*0202 | 3.1006% | Included |
| | DPA1*0301 | 0.0121% | Not included |
| | DPA1*0401 | 0.0009% | Not included |

Included: a term for the HLA allele was included in the empirical analysis; Baseline: the most frequent allele at each HLA locus, which was coded as the baseline allele in the analysis; Others: HLA alleles with a frequency lower than 0.5% that were merged into one group for the analysis; Not included: Rare alleles were not included in the analysis when the total frequency of grouped alleles at the locus was lower than 0.5%.

Table S6: Results from the two degree of freedom test of association between classical HLA alleles and birth weight in the UK Biobank, with and without adjustment for genetic variants in the HLA region (rs9366778 near HLA-C and rs6911024 in MICA) that have previously been associated with birth weight.

| **Locus** | **Allele** | **P value** | **P value (Conditioned on SNPs rs9366778 and rs6911024 )** |
|---|---|---|---|
| A | rs9366778* | NA | 2.02E-05 |
| | rs6911024* | NA | 9.05E-07 |
| | A*0101 | 1.09E-01 | 6.04E-02 |
| | **A*1101** | **8.14E-03** | 2.29E-01 |
| | A*0205 | 4.06E-01 | 4.73E-01 |
| | **A*2301** | **2.07E-03** | **6.20E-03** |
| | **A*2402** | **6.94E-03** | **3.33E-02** |
| | A*2501 | 7.17E-01 | 6.15E-01 |
| | **A*2601** | **3.23E-02** | 6.58E-02 |

| | | | |
|---|---|---|---|
| | **A*2902** | **3.86E-02** | 1.63E-01 |
| | A*3001 | 8.13E-02 | 5.57E-02 |
| | A*3002 | 6.45E-02 | 6.00E-02 |
| | **A*0301** | **7.39E-08** | **1.05E-06** |
| | **A*3101** | **1.03E-02** | **3.49E-02** |
| | A*3201 | 2.87E-01 | 2.86E-01 |
| | A*3301 | 7.38E-01 | 7.23E-01 |
| | A*6801 | 8.85E-02 | 2.44E-01 |
| | A*6802 | 4.62E-01 | 6.05E-01 |
| | A*Others | 4.45E-01 | 6.47E-01 |
| B | rs9366778 | NA | 1.08E-02 |
| | rs6911024 | NA | 6.11E-01 |
| | B*1302 | 7.81E-01 | 7.89E-01 |
| | B*1401 | 4.06E-01 | 4.12E-01 |
| | B*1402 | 8.50E-01 | 8.42E-01 |
| | B*1501 | 7.02E-01 | 1.75E-01 |
| | B*1801 | 6.89E-01 | 6.02E-01 |
| | B*2705 | 1.03E-01 | 1.07E-01 |
| | **B*3501** | **2.45E-09** | **3.24E-03** |
| | **B*3503** | **4.10E-02** | 2.17E-01 |
| | B*3701 | 5.44E-01 | 5.45E-01 |
| | **B*3801** | **2.26E-02** | **2.25E-02** |
| | B*3901 | 9.93E-01 | 8.26E-01 |
| | **B*3906** | **4.45E-04** | **8.50E-04** |
| | B*4001 | 5.51E-02 | 7.07E-01 |
| | B*4002 | 2.66E-01 | 2.74E-01 |
| | B*4402 | 8.50E-02 | **4.49E-02** |
| | **B*4403** | **3.28E-02** | 1.50E-01 |

| | | | |
|---|---|---|---|
| | B*4501 | 9.06E-01 | 9.20E-01 |
| | **B*4901** | **3.08E-02** | 2.83E-01 |
| | B*5001 | 6.35E-01 | 6.15E-01 |
| | **B*5101** | **1.38E-02** | 4.20E-01 |
| | B*5201 | 5.68E-01 | 4.35E-01 |
| | B*5501 | 4.10E-01 | 2.11E-01 |
| | B*5701 | 7.87E-01 | 7.79E-01 |
| | B*0801 | 5.82E-01 | 5.96E-01 |
| | B*Others | 1.24E-01 | 6.81E-02 |
| C | rs9366778 | NA | 1.88E-01 |
| | rs6911024 | NA | 2.81E-03 |
| | C*0102 | 1.97E-01 | 2.46E-01 |
| | C*1203 | 7.41E-01 | 7.93E-01 |
| | C*1402 | 8.15E-01 | 8.91E-01 |
| | **C*1502** | **4.26E-02** | 3.42E-01 |
| | C*1601 | 4.43E-01 | 4.08E-01 |
| | C*1701 | 8.28E-01 | 8.41E-01 |
| | C*0202 | 1.85E-01 | 1.89E-01 |
| | **C*0303** | **4.53E-03** | 5.14E-02 |
| | C*0304 | 6.84E-02 | 8.54E-01 |
| | **C*0401** | **2.13E-07** | 2.21E-01 |
| | **C*0501** | **6.91E-05** | **4.87E-04** |
| | C*0602 | 7.57E-01 | 8.70E-01 |
| | C*0702 | 6.15E-01 | 6.75E-01 |
| | C*0704 | 5.07E-01 | 9.82E-01 |
| | C*0802 | 7.71E-01 | 8.61E-01 |
| | C*Others | 3.57E-01 | 3.81E-01 |
| DPA1 | rs9366778 | NA | 2.11E-07 |

| | | | |
|---|---|---|---|
| | rs6911024 | NA | 3.00E-08 |
| | DPA1*0104 | 2.89E-01 | 2.89E-01 |
| | DPA1*0201 | 3.30E-01 | 2.20E-01 |
| | DPA1*0202 | 2.01E-01 | 2.36E-01 |
| DPB1 | rs9366778 | NA | 1.39E-06 |
| | rs6911024 | NA | 7.99E-08 |
| | DPB1*1001 | 8.82E-01 | 9.67E-01 |
| | **DPB1*0101** | **5.49E-04** | **1.15E-04** |
| | DPB1*1101 | 2.73E-01 | 2.01E-01 |
| | DPB1*1301 | 3.07E-01 | 3.29E-01 |
| | DPB1*1401 | 4.69E-01 | 4.07E-01 |
| | DPB1*1501 | 6.30E-02 | 6.35E-02 |
| | DPB1*1701 | 4.89E-01 | 4.89E-01 |
| | DPB1*1901 | 3.88E-01 | 4.82E-01 |
| | DPB1*0201 | 9.26E-01 | 9.75E-01 |
| | DPB1*0202 | 7.03E-01 | 8.22E-01 |
| | DPB1*0301 | 7.04E-02 | 9.59E-02 |
| | DPB1*0402 | 1.30E-01 | 3.59E-01 |
| | DPB1*0501 | 4.03E-01 | 3.60E-01 |
| | DPB1*0601 | 7.62E-02 | 1.34E-01 |
| | DPB1*0901 | 5.70E-01 | 5.66E-01 |
| | DPB1*Others | 1.84E-01 | 1.90E-01 |
| DQA1 | rs9366778 | NA | 9.77E-08 |
| | rs6911024 | NA | 4.45E-07 |
| | **DQA1*0101** | **6.22E-03** | 1.71E-01 |
| | DQA1*0102 | 7.92E-01 | 7.24E-01 |
| | DQA1*0103 | 7.40E-01 | 2.63E-01 |
| | DQA1*0201 | 7.53E-02 | 1.49E-01 |

| | | | |
|---|---|---|---|
| | DQA1*0301 | 7.10E-01 | 5.58E-01 |
| | DQA1*0401 | 1.74E-01 | 4.43E-01 |
| | DQA1*Others | 2.26E-01 | 1.55E-01 |
| DQB1 | rs9366778 | NA | 5.90E-06 |
| | rs6911024 | NA | 2.18E-06 |
| | DQB1*0201 | 8.08E-02 | 3.19E-01 |
| | DQB1*0202 | 1.38E-01 | 1.31E-01 |
| | DQB1*0302 | 1.86E-01 | 1.16E-01 |
| | DQB1*0303 | 6.06E-01 | 6.40E-01 |
| | DQB1*0402 | 1.95E-01 | 3.47E-01 |
| | **DQB1*0501** | **6.96E-03** | 7.87E-02 |
| | DQB1*0502 | 2.45E-01 | 3.87E-01 |
| | DQB1*0503 | 3.25E-01 | 4.87E-01 |
| | DQB1*0602 | 2.21E-01 | 6.14E-01 |
| | DQB1*0603 | 4.02E-01 | 2.91E-01 |
| | DQB1*0604 | 5.22E-01 | 5.84E-01 |
| | DQB1*0609 | 5.10E-02 | 6.77E-02 |
| | DQB1*Others | 6.01E-01 | 4.05E-01 |
| DRB1 | rs9366778 | NA | 2.05E-05 |
| | rs6911024 | NA | 8.71E-06 |
| | DRB1*1001 | 3.85E-01 | 4.43E-01 |
| | **DRB1*0101** | **1.52E-02** | 3.74E-01 |
| | DRB1*0102 | 5.05E-01 | 4.66E-01 |
| | **DRB1*0103** | **1.75E-02** | 1.77E-01 |
| | DRB1*1101 | 3.04E-01 | 7.33E-01 |
| | **DRB1*1104** | **3.60E-03** | **1.80E-02** |
| | DRB1*1201 | 3.21E-01 | 3.97E-01 |
| | DRB1*1301 | 7.95E-01 | 6.04E-01 |

| | | | |
|---|---|---|---|
| | DRB1*1302 | 2.30E-01 | 7.89E-01 |
| | DRB1*1303 | 2.89E-01 | 3.68E-01 |
| | DRB1*1401 | 5.30E-01 | 7.81E-01 |
| | DRB1*1501 | 5.39E-01 | 5.49E-01 |
| | DRB1*1601 | 6.96E-01 | 9.35E-01 |
| | DRB1*0401 | 3.98E-01 | 1.91E-01 |
| | **DRB1*0404** | **1.56E-02** | 2.08E-01 |
| | DRB1*0407 | 8.84E-02 | 3.31E-01 |
| | **DRB1*0701** | **4.78E-02** | 2.87E-01 |
| | **DRB1*0801** | **1.74E-02** | 7.91E-02 |
| | DRB1*0901 | 6.51E-01 | 9.31E-01 |
| | DRB1*Others | 3.20E-01 | 4.65E-01 |
| DRB3 | rs9366778 | NA | 6.43E-07 |
| | rs6911024 | NA | 7.40E-08 |
| | DRB3*0101 | 1.73E-01 | 8.67E-01 |
| | DRB3*0202 | 2.42E-01 | 1.94E-01 |
| | DRB3*0301 | 6.98E-01 | 8.18E-01 |
| DRB4 | rs9366778 | NA | 4.43E-06 |
| | rs6911024 | NA | 3.49E-08 |
| | **DRB4*0101** | **2.03E-02** | **4.34E-02** |
| | DRB4*0103 | 9.64E-01 | 9.29E-01 |
| DRB5 | rs9366778 | NA | 3.04E-07 |
| | rs6911024 | NA | 1.51E-07 |
| | **DRB5*0101** | **2.35E-02** | 2.20E-01 |
| | DRB5*0202 | 4.92E-01 | 7.32E-01 |

The bold font indicates the alleles with a significance level the P < 0.05, The pink highlight indicates the alleles with a significance level the P < 0.001; * Effect alleles: rs9366778-A and rs6911024-C.

Table S7: Results from the structural equation model for 19 HLA alleles that had nominally significant (P<0.05) maternal and/or fetal effects on birth weight in the UK Biobank.

| HLA loci | Allele | Frequency | Maternal Effect Size | Maternal SE | Maternal P-Value | Fetal Effect Size | Fetal SE | Fetal P-Value |
|---|---|---|---|---|---|---|---|---|
| A | A*0101 | 19.45% | -0.0147 | 0.0071 | 3.88E-02 | 0.0087 | 0.0068 | 2.04E-01 |
| | A*0301 | 14.51% | -0.0419 | 0.0078 | 7.90E-08 | 0.0202 | 0.0075 | 6.85E-03 |
| | A*2301 | 1.69% | -0.0425 | 0.0191 | 2.61E-02 | -0.0020 | 0.0183 | 9.13E-01 |
| | A*2402 | 7.37% | -0.0260 | 0.0100 | 9.23E-03 | 0.0075 | 0.0096 | 4.33E-01 |
| B | B*3501 | 4.68% | -0.0516 | 0.0131 | 7.78E-05 | -0.0019 | 0.0125 | 8.80E-01 |
| | B*3801 | 0.76% | -0.0768 | 0.0277 | 5.60E-03 | 0.0449 | 0.0265 | 9.03E-02 |
| | B*3906 | 0.69% | -0.1133 | 0.0288 | 8.49E-05 | 0.0986 | 0.0279 | 4.03E-04 |
| C | C*0303 | 5.57% | 0.0379 | 0.0117 | 1.26E-03 | -0.0228 | 0.0112 | 4.21E-02 |
| | C*0401 | 8.32% | -0.0272 | 0.0101 | 7.20E-03 | -0.0113 | 0.0097 | 2.47E-01 |
| DRB1 | DRB1*0101 | 9.35% | -0.0221 | 0.0102 | 2.94E-02 | 0.0039 | 0.0098 | 6.88E-01 |
| | DRB1*0801 | 1.82% | 0.0109 | 0.0193 | 5.72E-01 | -0.0415 | 0.0184 | 2.44E-02 |
| | DRB1*1104 | 1.16% | 0.0440 | 0.0236 | 6.17E-02 | -0.0727 | 0.0225 | 1.21E-03 |
| DQB1 | DQB1*0201 | 15.00% | -0.0107 | 0.0084 | 2.04E-01 | 0.0176 | 0.0081 | 2.94E-02 |
| | DQB1*0501 | 12.01% | -0.0276 | 0.0090 | 2.10E-03 | 0.0159 | 0.0086 | 6.46E-02 |
| | DQB1*0609 | 1.04% | -0.0587 | 0.0243 | 1.57E-02 | 0.0356 | 0.0234 | 1.28E-01 |
| DQA1 | DQA1*0101 | 14.14% | -0.0227 | 0.0081 | 5.05E-03 | 0.0087 | 0.0078 | 2.63E-01 |
| DPB1 | DPB1*0101 | 5.91% | -0.0255 | 0.0107 | 1.72E-02 | -0.0022 | 0.0103 | 8.29E-01 |
| | DPB1*0301 | 10.33% | -0.0185 | 0.0084 | 2.79E-02 | 0.0097 | 0.0081 | 2.30E-01 |
| | DPB1*1501 | 0.74% | 0.0380 | 0.0289 | 1.89E-01 | -0.0623 | 0.0274 | 2.33E-02 |

SE: standard error

Table S8: Results from the structural equation model for all HLA alleles with estimated maternal and fetal effects on birth weight in the UK Biobank, with and without adjustment for genetic variants in the HLA region (rs9366778 near HLA-C and rs6911024 in MICA) that have previously been associated with birth weight.

| | | | Unconditional analysis | | | | | | Conditioned on SNPs rs9366778 and rs6911024 | | | | | |
|---|---|---|---|---|---|---|---|---|---|---|---|---|---|---|
| | | | Maternal effect | | | Fetal effect | | | Maternal effect | | | Fetal effect | | |
| Locus | Allele/SNP | Frequency | Estimated effect sizes | SE | P value | Estimated effect sizes | SE | P value | Estimated effect sizes | SE | P value | Estimated effect sizes | SE | P value |
| | rs9366778[1] | NA | NA | NA | NA | NA | NA | NA | 0.0047 | 0.0056 | 3.96E-01 | -0.0194 | 0.0053 | 2.67E-04 |
| | rs6911024[1] | NA | NA | NA | NA | NA | NA | NA | -0.0386 | 0.0089 | 1.34E-05 | 0.0112 | 0.0085 | 1.88E-01 |
| A | A*0101 | 19.45% | -0.0147 | 0.0071 | 3.88E-02 | 0.0087 | 0.0068 | 2.04E-01 | -0.0143 | 0.0073 | 4.92E-02 | 0.0043 | 0.0070 | 5.34E-01 |
| A | A*0205 | 0.75% | -0.0259 | 0.0280 | 3.55E-01 | 0.0014 | 0.0268 | 9.57E-01 | -0.0238 | 0.0280 | 3.95E-01 | 0.0015 | 0.0268 | 9.54E-01 |
| A | A*0301 | 14.51% | -0.0419 | 0.0078 | 7.90E-08 | 0.0202 | 0.0075 | 6.85E-03 | -0.0383 | 0.0079 | 1.17E-06 | 0.0181 | 0.0076 | 1.71E-02 |
| A | A*1101 | 6.06% | -0.0203 | 0.0108 | 5.98E-02 | -0.0021 | 0.0104 | 8.37E-01 | -0.0129 | 0.0110 | 2.43E-01 | 0.0006 | 0.0106 | 9.57E-01 |
| A | A*2301 | 1.69% | -0.0425 | 0.0191 | 2.61E-02 | -0.0020 | 0.0183 | 9.13E-01 | -0.0450 | 0.0192 | 1.94E-02 | 0.0062 | 0.0185 | 7.36E-01 |
| A | A*2402 | 7.37% | -0.0260 | 0.0100 | 9.23E-03 | 0.0075 | 0.0096 | 4.33E-01 | -0.0223 | 0.0101 | 2.67E-02 | 0.0075 | 0.0097 | 4.38E-01 |
| A | A*2501 | 1.73% | 0.0102 | 0.0189 | 5.88E-01 | -0.0001 | 0.0181 | 9.94E-01 | 0.0157 | 0.0190 | 4.10E-01 | -0.0049 | 0.0183 | 7.90E-01 |
| A | A*2601 | 1.97% | -0.0281 | 0.0177 | 1.13E-01 | -0.0031 | 0.0171 | 8.55E-01 | -0.0225 | 0.0179 | 2.10E-01 | -0.0058 | 0.0173 | 7.35E-01 |
| A | A*2902 | 4.24% | -0.0106 | 0.0125 | 3.97E-01 | -0.0114 | 0.0120 | 3.42E-01 | -0.0134 | 0.0128 | 2.94E-01 | -0.0031 | 0.0123 | 8.00E-01 |
| A | A*3001 | 1.00% | -0.0448 | 0.0244 | 6.59E-02 | 0.0128 | 0.0235 | 5.85E-01 | -0.0445 | 0.0244 | 6.85E-02 | 0.0081 | 0.0235 | 7.31E-01 |
| A | A*3002 | 0.98% | -0.0484 | 0.0248 | 5.10E-02 | 0.0149 | 0.0238 | 5.33E-01 | -0.0476 | 0.0249 | 5.56E-02 | 0.0126 | 0.0238 | 5.96E-01 |
| A | A*3101 | 2.65% | -0.0237 | 0.0155 | 1.27E-01 | -0.0084 | 0.0148 | 5.72E-01 | -0.0240 | 0.0156 | 1.23E-01 | -0.0031 | 0.0149 | 8.32E-01 |
| A | A*3201 | 3.46% | -0.0150 | 0.0138 | 2.76E-01 | 0.0009 | 0.0132 | 9.46E-01 | -0.0142 | 0.0138 | 3.03E-01 | -0.0002 | 0.0132 | 9.91E-01 |
| A | A*3301 | 0.54% | 0.0256 | 0.0330 | 4.37E-01 | -0.0176 | 0.0317 | 5.80E-01 | 0.0258 | 0.0325 | 4.28E-01 | -0.0227 | 0.0311 | 4.66E-01 |
| A | A*6801 | 3.00% | -0.0018 | 0.0145 | 9.00E-01 | -0.0190 | 0.0140 | 1.76E-01 | -0.0029 | 0.0147 | 8.45E-01 | -0.0134 | 0.0142 | 3.43E-01 |
| A | A*6802 | 0.59% | -0.0380 | 0.0314 | 2.26E-01 | 0.0218 | 0.0304 | 4.73E-01 | -0.0310 | 0.0315 | 3.24E-01 | 0.0188 | 0.0304 | 5.37E-01 |
| A | A*Other | 1.65% | -0.0168 | 0.0178 | 3.46E-01 | 0.0025 | 0.0170 | 8.85E-01 | -0.0128 | 0.0179 | 4.74E-01 | 0.0025 | 0.0171 | 8.84E-01 |

| | | | | | | | | | | | | | | |
|---|---|---|---|---|---|---|---|---|---|---|---|---|---|---|
| | rs9366778 | NA | NA | NA | NA | NA | NA | NA | 0.0137 | 0.0114 | 2.29E-01 | -0.0292 | 0.0106 | 5.74E-03 |
| | rs6911024 | NA | NA | NA | NA | NA | NA | NA | 0.0205 | 0.0205 | 3.17E-01 | -0.0186 | 0.0198 | 3.48E-01 |
| | B*0801 | 14.45% | -0.0090 | 0.0092 | 3.25E-01 | 0.0080 | 0.0087 | 3.59E-01 | -0.0089 | 0.0096 | 3.58E-01 | 0.0077 | 0.0087 | 3.74E-01 |
| | B*1302 | 1.92% | -0.0025 | 0.0186 | 8.92E-01 | 0.0099 | 0.0179 | 5.80E-01 | -0.0023 | 0.0188 | 9.01E-01 | 0.0096 | 0.0177 | 5.87E-01 |
| | B*1401 | 1.19% | -0.0288 | 0.0230 | 2.10E-01 | 0.0283 | 0.0219 | 1.98E-01 | -0.0286 | 0.0234 | 2.20E-01 | 0.0279 | 0.0219 | 2.03E-01 |
| | B*1402 | 2.55% | -0.0029 | 0.0165 | 8.61E-01 | -0.0035 | 0.0158 | 8.23E-01 | -0.0027 | 0.0170 | 8.72E-01 | -0.0038 | 0.0159 | 8.09E-01 |
| | B*1501 | 6.21% | 0.0097 | 0.0118 | 4.11E-01 | -0.0074 | 0.0112 | 5.08E-01 | -0.0037 | 0.0172 | 8.28E-01 | 0.0213 | 0.0153 | 1.65E-01 |
| | B*1801 | 3.67% | 0.0120 | 0.0143 | 4.04E-01 | -0.0072 | 0.0136 | 5.97E-01 | 0.0070 | 0.0155 | 6.50E-01 | 0.0031 | 0.0141 | 8.27E-01 |
| | B*2705 | 3.95% | 0.0014 | 0.0139 | 9.18E-01 | -0.0194 | 0.0133 | 1.42E-01 | 0.0013 | 0.0142 | 9.29E-01 | -0.0192 | 0.0132 | 1.46E-01 |
| | B*3501 | 4.68% | -0.0516 | 0.0131 | 7.78E-05 | -0.0019 | 0.0125 | 8.80E-01 | -0.0854 | 0.0274 | 1.83E-03 | 0.0452 | 0.0246 | 6.66E-02 |
| | B*3503 | 0.85% | -0.0125 | 0.0268 | 6.41E-01 | -0.0333 | 0.0256 | 1.94E-01 | -0.0308 | 0.0306 | 3.15E-01 | -0.0035 | 0.0281 | 9.00E-01 |
| | B*3701 | 1.38% | -0.0231 | 0.0215 | 2.83E-01 | 0.0130 | 0.0205 | 5.25E-01 | -0.0231 | 0.0220 | 2.93E-01 | 0.0130 | 0.0205 | 5.27E-01 |
| | B*3801 | 0.76% | -0.0768 | 0.0277 | 5.60E-03 | 0.0449 | 0.0265 | 9.03E-02 | -0.0971 | 0.0355 | 6.21E-03 | 0.0636 | 0.0333 | 5.63E-02 |
| B | B*3901 | 0.78% | -0.0016 | 0.0280 | 9.55E-01 | -0.0007 | 0.0268 | 9.78E-01 | -0.0217 | 0.0345 | 5.29E-01 | 0.0175 | 0.0322 | 5.86E-01 |
| | B*3906 | 0.69% | -0.1133 | 0.0288 | 8.49E-05 | 0.0986 | 0.0279 | 4.03E-04 | -0.1336 | 0.0355 | 1.65E-04 | 0.1169 | 0.0337 | 5.31E-04 |
| | B*4001 | 5.52% | -0.0001 | 0.0122 | 9.96E-01 | -0.0183 | 0.0116 | 1.16E-01 | -0.0135 | 0.0177 | 4.46E-01 | 0.0105 | 0.0157 | 5.06E-01 |
| | B*4002 | 0.83% | -0.0310 | 0.0271 | 2.53E-01 | 0.0026 | 0.0262 | 9.22E-01 | -0.0317 | 0.0268 | 2.38E-01 | 0.0040 | 0.0256 | 8.77E-01 |
| | B*4402 | 11.22% | 0.0082 | 0.0097 | 3.99E-01 | 0.0064 | 0.0093 | 4.91E-01 | 0.0066 | 0.0104 | 5.24E-01 | 0.0098 | 0.0093 | 2.95E-01 |
| | B*4403 | 5.89% | -0.0175 | 0.0120 | 1.43E-01 | -0.0033 | 0.0114 | 7.70E-01 | -0.0308 | 0.0173 | 7.42E-02 | 0.0251 | 0.0154 | 1.03E-01 |
| | B*4501 | 0.65% | 0.0042 | 0.0297 | 8.86E-01 | -0.0113 | 0.0288 | 6.94E-01 | 0.0038 | 0.0306 | 9.02E-01 | -0.0104 | 0.0293 | 7.23E-01 |
| | B*4901 | 1.14% | -0.0132 | 0.0237 | 5.76E-01 | -0.0288 | 0.0227 | 2.05E-01 | -0.0265 | 0.0267 | 3.22E-01 | -0.0006 | 0.0245 | 9.80E-01 |
| | B*5001 | 0.85% | -0.0247 | 0.0269 | 3.59E-01 | 0.0225 | 0.0256 | 3.80E-01 | -0.0252 | 0.0268 | 3.46E-01 | 0.0237 | 0.0252 | 3.48E-01 |
| | B*5101 | 3.66% | 0.0007 | 0.0143 | 9.62E-01 | -0.0266 | 0.0137 | 5.17E-02 | -0.0093 | 0.0173 | 5.93E-01 | -0.0055 | 0.0157 | 7.27E-01 |
| | B*5201 | 0.51% | -0.0320 | 0.0343 | 3.51E-01 | 0.0116 | 0.0329 | 7.24E-01 | -0.0455 | 0.0358 | 2.05E-01 | 0.0402 | 0.0334 | 2.29E-01 |
| | B*5501 | 1.83% | 0.0246 | 0.0191 | 1.97E-01 | -0.0136 | 0.0182 | 4.55E-01 | 0.0111 | 0.0229 | 6.27E-01 | 0.0154 | 0.0209 | 4.63E-01 |
| | B*5701 | 3.92% | -0.0092 | 0.0139 | 5.05E-01 | 0.0053 | 0.0133 | 6.92E-01 | -0.0096 | 0.0144 | 5.05E-01 | 0.0060 | 0.0133 | 6.51E-01 |
| | B*Other | 3.62% | -0.0218 | 0.0134 | 1.03E-01 | 0.0058 | 0.0128 | 6.48E-01 | -0.0369 | 0.0175 | 3.47E-02 | 0.0303 | 0.0155 | 5.06E-02 |
| C | rs9366778 | NA | NA | NA | NA | NA | NA | NA | 0.0103 | 0.0126 | 4.14E-01 | -0.0206 | 0.0121 | 8.90E-02 |

| | | | | | | | | | | | | | | |
|---|---|---|---|---|---|---|---|---|---|---|---|---|---|---|
| | rs6911024 | NA | NA | NA | NA | NA | NA | NA | -0.0377 | 0.0115 | 1.02E-03 | 0.0202 | 0.0111 | 6.77E-02 |
| | C*0102 | 3.40% | 0.0109 | 0.0143 | 4.44E-01 | -0.0226 | 0.0137 | 9.91E-02 | 0.0100 | 0.0143 | 4.85E-01 | -0.0210 | 0.0137 | 1.27E-01 |
| | C*0202 | 3.60% | -0.0241 | 0.0139 | 8.35E-02 | 0.0120 | 0.0134 | 3.68E-01 | -0.0226 | 0.0141 | 1.09E-01 | 0.0086 | 0.0135 | 5.24E-01 |
| | C*0303 | 5.57% | 0.0379 | 0.0117 | 1.26E-03 | -0.0228 | 0.0112 | 4.21E-02 | 0.0294 | 0.0155 | 5.86E-02 | -0.0062 | 0.0148 | 6.74E-01 |
| | C*0304 | 7.96% | 0.0042 | 0.0103 | 6.86E-01 | -0.0178 | 0.0099 | 7.18E-02 | -0.0049 | 0.0146 | 7.36E-01 | -0.0006 | 0.0140 | 9.67E-01 |
| | C*0401 | 8.32% | -0.0272 | 0.0101 | 7.20E-03 | -0.0113 | 0.0097 | 2.47E-01 | -0.0115 | 0.0158 | 4.65E-01 | -0.0074 | 0.0151 | 6.26E-01 |
| | C*0501 | 11.30% | 0.0143 | 0.0092 | 1.18E-01 | 0.0133 | 0.0088 | 1.32E-01 | 0.0155 | 0.0095 | 1.01E-01 | 0.0099 | 0.0091 | 2.76E-01 |
| | C*0602 | 8.97% | -0.0066 | 0.0099 | 5.04E-01 | 0.0068 | 0.0095 | 4.72E-01 | -0.0053 | 0.0101 | 6.00E-01 | 0.0035 | 0.0097 | 7.21E-01 |
| | C*0702 | 15.93% | 0.0028 | 0.0083 | 7.39E-01 | 0.0029 | 0.0080 | 7.22E-01 | 0.0062 | 0.0087 | 4.75E-01 | -0.0016 | 0.0083 | 8.45E-01 |
| | C*0704 | 1.75% | 0.0081 | 0.0190 | 6.69E-01 | -0.0190 | 0.0183 | 3.01E-01 | -0.0012 | 0.0215 | 9.57E-01 | -0.0016 | 0.0207 | 9.37E-01 |
| | C*0802 | 3.67% | -0.0086 | 0.0138 | 5.34E-01 | 0.0094 | 0.0133 | 4.81E-01 | -0.0076 | 0.0140 | 5.88E-01 | 0.0061 | 0.0135 | 6.50E-01 |
| | C*1203 | 2.89% | -0.0116 | 0.0153 | 4.50E-01 | 0.0099 | 0.0147 | 4.99E-01 | 0.0102 | 0.0167 | 5.40E-01 | -0.0043 | 0.0160 | 7.87E-01 |
| | C*1402 | 0.91% | 0.0155 | 0.0258 | 5.47E-01 | -0.0147 | 0.0248 | 5.53E-01 | 0.0071 | 0.0277 | 7.98E-01 | 0.0020 | 0.0266 | 9.40E-01 |
| | C*1502 | 1.74% | 0.0020 | 0.0191 | 9.17E-01 | -0.0318 | 0.0184 | 8.40E-02 | -0.0065 | 0.0217 | 7.65E-01 | -0.0150 | 0.0209 | 4.72E-01 |
| | C*1601 | 4.46% | -0.0120 | 0.0128 | 3.48E-01 | 0.0017 | 0.0123 | 8.89E-01 | -0.0212 | 0.0164 | 1.96E-01 | 0.0190 | 0.0157 | 2.28E-01 |
| | C*1701 | 0.53% | 0.0158 | 0.0328 | 6.31E-01 | -0.0032 | 0.0318 | 9.20E-01 | 0.0173 | 0.0332 | 6.02E-01 | -0.0068 | 0.0321 | 8.33E-01 |
| | C*Other | 1.02% | -0.0320 | 0.0243 | 1.87E-01 | 0.0148 | 0.0232 | 5.25E-01 | -0.0360 | 0.0260 | 1.67E-01 | 0.0286 | 0.0250 | 2.52E-01 |
| | rs9366778 | NA | NA | NA | NA | NA | NA | NA | 0.0048 | 0.0051 | 3.53E-01 | -0.0211 | 0.0049 | 1.76E-05 |
| | rs6911024 | NA | NA | NA | NA | NA | NA | NA | -0.0424 | 0.0086 | 8.31E-07 | 0.0132 | 0.0083 | 1.11E-01 |
| DPA1 | DPA1*0104 | 14.14% | -0.0227 | 0.0081 | 5.05E-03 | 0.0087 | 0.0078 | 2.63E-01 | 0.0395 | 0.0335 | 2.38E-01 | -0.0504 | 0.0320 | 1.15E-01 |
| | DPA1*0201 | 19.31% | 0.0047 | 0.0074 | 5.25E-01 | -0.0023 | 0.0071 | 7.52E-01 | -0.0095 | 0.0068 | 1.63E-01 | 0.0024 | 0.0066 | 7.12E-01 |
| | DPA1*0202 | 5.49% | 0.0006 | 0.0114 | 9.62E-01 | 0.0052 | 0.0110 | 6.37E-01 | -0.0080 | 0.0139 | 5.63E-01 | -0.0082 | 0.0134 | 5.38E-01 |
| | rs9366778 | 14.66% | -0.0107 | 0.0080 | 1.82E-01 | -0.0016 | 0.0077 | 8.37E-01 | 0.0038 | 0.0054 | 4.84E-01 | -0.0203 | 0.0052 | 9.10E-05 |
| | rs6911024 | 20.25% | -0.0049 | 0.0073 | 5.03E-01 | 0.0013 | 0.0070 | 8.54E-01 | -0.0430 | 0.0090 | 1.59E-06 | 0.0136 | 0.0086 | 1.16E-01 |
| DPB1 | DPB1*0101 | 1.94% | 0.0012 | 0.0180 | 9.46E-01 | -0.0221 | 0.0173 | 2.01E-01 | -0.0270 | 0.0108 | 1.20E-02 | -0.0038 | 0.0103 | 7.15E-01 |
| | DPB1*0201 | 0.51% | 0.0434 | 0.0297 | 1.43E-01 | -0.0146 | 0.0287 | 6.12E-01 | -0.0015 | 0.0082 | 8.59E-01 | 0.0018 | 0.0079 | 8.21E-01 |
| | DPB1*0202 | NA | NA | NA | NA | NA | NA | NA | 0.0070 | 0.0282 | 8.05E-01 | 0.0053 | 0.0273 | 8.47E-01 |

| | | | | | | | | | | | | | | |
|---|---|---|---|---|---|---|---|---|---|---|---|---|---|---|
| | DPB1*0301 | NA | NA | NA | NA | NA | NA | NA | -0.0180 | 0.0084 | 3.23E-02 | 0.0113 | 0.0081 | 1.62E-01 |
| | DPB1*0402 | 15.00% | -0.0107 | 0.0084 | 2.04E-01 | 0.0176 | 0.0081 | 2.94E-02 | -0.0034 | 0.0083 | 6.80E-01 | -0.0047 | 0.0080 | 5.55E-01 |
| | DPB1*0501 | 10.45% | -0.0134 | 0.0094 | 1.55E-01 | 0.0014 | 0.0090 | 8.79E-01 | -0.0201 | 0.0173 | 2.46E-01 | 0.0054 | 0.0167 | 7.48E-01 |
| | DPB1*0601 | 10.27% | -0.0170 | 0.0095 | 7.21E-02 | 0.0101 | 0.0091 | 2.67E-01 | -0.0313 | 0.0226 | 1.66E-01 | 0.0018 | 0.0218 | 9.36E-01 |
| | DPB1*0901 | 5.23% | -0.0105 | 0.0120 | 3.79E-01 | 0.0113 | 0.0115 | 3.27E-01 | 0.0126 | 0.0339 | 7.10E-01 | 0.0123 | 0.0326 | 7.07E-01 |
| | DPB1*1001 | 1.99% | -0.0080 | 0.0180 | 6.56E-01 | -0.0141 | 0.0173 | 4.14E-01 | 0.0005 | 0.0194 | 9.80E-01 | -0.0035 | 0.0187 | 8.51E-01 |
| | DPB1*1101 | 12.01% | -0.0276 | 0.0090 | 2.10E-03 | 0.0159 | 0.0086 | 6.46E-02 | -0.0277 | 0.0157 | 7.74E-02 | 0.0230 | 0.0150 | 1.26E-01 |
| | DPB1*1301 | 0.69% | -0.0363 | 0.0293 | 2.15E-01 | 0.0050 | 0.0280 | 8.60E-01 | -0.0167 | 0.0204 | 4.15E-01 | -0.0040 | 0.0197 | 8.37E-01 |
| | DPB1*1401 | 2.15% | -0.0251 | 0.0175 | 1.52E-01 | 0.0131 | 0.0167 | 4.32E-01 | 0.0145 | 0.0258 | 5.74E-01 | 0.0094 | 0.0248 | 7.04E-01 |
| | DPB1*1501 | 14.42% | 0.0013 | 0.0085 | 8.83E-01 | 0.0084 | 0.0082 | 3.02E-01 | 0.0368 | 0.0289 | 2.02E-01 | -0.0619 | 0.0274 | 2.38E-02 |
| | DPB1*1701 | 5.16% | -0.0054 | 0.0120 | 6.51E-01 | 0.0136 | 0.0115 | 2.38E-01 | -0.0159 | 0.0234 | 4.95E-01 | -0.0030 | 0.0225 | 8.94E-01 |
| | DPB1*1901 | 2.75% | 0.0171 | 0.0155 | 2.72E-01 | -0.0155 | 0.0149 | 2.99E-01 | 0.0098 | 0.0336 | 7.71E-01 | -0.0324 | 0.0326 | 3.20E-01 |
| | DPB1*Other | 1.04% | -0.0587 | 0.0243 | 1.57E-02 | 0.0356 | 0.0234 | 1.28E-01 | -0.0419 | 0.0258 | 1.04E-01 | 0.0439 | 0.0249 | 7.75E-02 |
| | rs9366778 | 0.70% | -0.0171 | 0.0291 | 5.56E-01 | 0.0276 | 0.0279 | 3.22E-01 | 0.0070 | 0.0053 | 1.84E-01 | -0.0235 | 0.0051 | 3.58E-06 |
| | rs6911024 | NA | NA | NA | NA | NA | NA | NA | -0.0405 | 0.0090 | 6.45E-06 | 0.0124 | 0.0086 | 1.52E-01 |
| | DQA1*0101 | NA | NA | NA | NA | NA | NA | NA | -0.0157 | 0.0084 | 6.23E-02 | 0.0126 | 0.0081 | 1.18E-01 |
| | DQA1*0102 | 0.52% | 0.0408 | 0.0336 | 2.25E-01 | -0.0503 | 0.0320 | 1.16E-01 | 0.0045 | 0.0074 | 5.48E-01 | -0.0007 | 0.0071 | 9.18E-01 |
| DQA1 | DQA1*0103 | 14.60% | -0.0085 | 0.0068 | 2.13E-01 | 0.0026 | 0.0066 | 6.87E-01 | 0.0015 | 0.0115 | 8.94E-01 | 0.0107 | 0.0111 | 3.31E-01 |
| | DQA1*0201 | 3.10% | -0.0068 | 0.0139 | 6.25E-01 | -0.0102 | 0.0134 | 4.44E-01 | -0.0130 | 0.0081 | 1.10E-01 | 0.0036 | 0.0078 | 6.39E-01 |
| | DQA1*0301 | NA | NA | NA | NA | NA | NA | NA | -0.0068 | 0.0074 | 3.60E-01 | 0.0076 | 0.0071 | 2.87E-01 |
| | DQA1*0401 | NA | NA | NA | NA | NA | NA | NA | 0.0068 | 0.0181 | 7.06E-01 | -0.0188 | 0.0173 | 2.77E-01 |
| | DQA1*Other | 5.91% | -0.0255 | 0.0107 | 1.72E-02 | -0.0022 | 0.0103 | 8.29E-01 | 0.0443 | 0.0298 | 1.37E-01 | -0.0091 | 0.0288 | 7.51E-01 |
| | rs9366778 | 11.18% | -0.0024 | 0.0082 | 7.66E-01 | 0.0005 | 0.0079 | 9.54E-01 | 0.0063 | 0.0056 | 2.59E-01 | -0.0214 | 0.0054 | 6.59E-05 |
| | rs6911024 | 0.76% | 0.0084 | 0.0282 | 7.65E-01 | 0.0080 | 0.0273 | 7.70E-01 | -0.0381 | 0.0090 | 2.55E-05 | 0.0110 | 0.0087 | 2.04E-01 |
| DQB1 | DQB1*0201 | 10.33% | -0.0185 | 0.0084 | 2.79E-02 | 0.0097 | 0.0081 | 2.30E-01 | -0.0113 | 0.0086 | 1.89E-01 | 0.0122 | 0.0083 | 1.39E-01 |
| | DQB1*0202 | 10.77% | -0.0060 | 0.0083 | 4.72E-01 | -0.0056 | 0.0080 | 4.83E-01 | -0.0163 | 0.0095 | 8.61E-02 | 0.0055 | 0.0091 | 5.47E-01 |
| | DQB1*0302 | 2.09% | -0.0196 | 0.0173 | 2.59E-01 | 0.0061 | 0.0167 | 7.15E-01 | -0.0197 | 0.0096 | 3.98E-02 | 0.0159 | 0.0092 | 8.37E-02 |

| | | | | | | | | | | | | | | |
|---|---|---|---|---|---|---|---|---|---|---|---|---|---|---|
| | DQB1*0303 | 1.21% | -0.0290 | 0.0226 | 1.99E-01 | -0.0058 | 0.0218 | 7.91E-01 | -0.0113 | 0.0120 | 3.47E-01 | 0.0089 | 0.0115 | 4.42E-01 |
| | DQB1*0402 | 0.55% | 0.0124 | 0.0341 | 7.16E-01 | 0.0122 | 0.0329 | 7.10E-01 | -0.0029 | 0.0182 | 8.72E-01 | -0.0144 | 0.0174 | 4.09E-01 |
| | DQB1*0501 | 1.66% | -0.0019 | 0.0194 | 9.22E-01 | -0.0046 | 0.0186 | 8.03E-01 | -0.0206 | 0.0092 | 2.49E-02 | 0.0162 | 0.0088 | 6.71E-02 |
| | DQB1*0502 | 2.63% | -0.0247 | 0.0155 | 1.12E-01 | 0.0154 | 0.0149 | 3.02E-01 | -0.0288 | 0.0299 | 3.36E-01 | 0.0025 | 0.0286 | 9.31E-01 |
| | DQB1*0503 | 1.49% | -0.0165 | 0.0205 | 4.20E-01 | -0.0049 | 0.0197 | 8.03E-01 | -0.0211 | 0.0176 | 2.30E-01 | 0.0154 | 0.0168 | 3.59E-01 |
| | DQB1*0602 | 0.92% | 0.0128 | 0.0257 | 6.19E-01 | 0.0091 | 0.0248 | 7.13E-01 | 0.0006 | 0.0086 | 9.44E-01 | 0.0049 | 0.0083 | 5.52E-01 |
| | DQB1*0603 | 0.74% | 0.0380 | 0.0289 | 1.89E-01 | -0.0623 | 0.0274 | 2.33E-02 | -0.0042 | 0.0121 | 7.31E-01 | 0.0147 | 0.0116 | 2.06E-01 |
| | DQB1*0604 | 1.12% | -0.0156 | 0.0234 | 5.06E-01 | -0.0034 | 0.0225 | 8.79E-01 | 0.0153 | 0.0158 | 3.32E-01 | -0.0075 | 0.0151 | 6.21E-01 |
| | DQB1*0609 | 0.52% | 0.0107 | 0.0338 | 7.51E-01 | -0.0367 | 0.0328 | 2.63E-01 | -0.0554 | 0.0247 | 2.51E-02 | 0.0319 | 0.0238 | 1.80E-01 |
| | DQB1*Other | 0.73% | -0.0437 | 0.0258 | 9.06E-02 | 0.0434 | 0.0249 | 8.20E-02 | -0.0212 | 0.0295 | 4.71E-01 | 0.0364 | 0.0283 | 1.98E-01 |
| | rs9366778 | NA | NA | NA | NA | NA | NA | NA | 0.0070 | 0.0056 | 2.14E-01 | -0.0208 | 0.0054 | 1.01E-04 |
| | rs6911024 | NA | NA | NA | NA | NA | NA | NA | -0.0391 | 0.0093 | 2.49E-05 | 0.0145 | 0.0089 | 1.04E-01 |
| | DRB1*0101 | 9.35% | -0.0221 | 0.0102 | 2.94E-02 | 0.0039 | 0.0098 | 6.88E-01 | -0.0147 | 0.0106 | 1.67E-01 | 0.0092 | 0.0102 | 3.67E-01 |
| | DRB1*0102 | 0.72% | 0.0132 | 0.0294 | 6.54E-01 | 0.0104 | 0.0282 | 7.12E-01 | 0.0134 | 0.0293 | 6.46E-01 | 0.0114 | 0.0280 | 6.82E-01 |
| | DRB1*0103 | 1.74% | -0.0066 | 0.0195 | 7.35E-01 | -0.0300 | 0.0188 | 1.11E-01 | 0.0051 | 0.0199 | 7.96E-01 | -0.0268 | 0.0192 | 1.61E-01 |
| | DRB1*0401 | 11.43% | 0.0106 | 0.0096 | 2.70E-01 | -0.0030 | 0.0092 | 7.47E-01 | 0.0086 | 0.0097 | 3.73E-01 | 0.0034 | 0.0092 | 7.09E-01 |
| | DRB1*0404 | 4.05% | -0.0010 | 0.0136 | 9.42E-01 | -0.0240 | 0.0131 | 6.58E-02 | -0.0033 | 0.0140 | 8.12E-01 | -0.0130 | 0.0134 | 3.30E-01 |
| | DRB1*0407 | 0.98% | -0.0302 | 0.0256 | 2.38E-01 | -0.0081 | 0.0245 | 7.41E-01 | -0.0263 | 0.0257 | 3.08E-01 | 0.0014 | 0.0246 | 9.55E-01 |
| DRB1 | DRB1*0701 | 14.64% | -0.0055 | 0.0090 | 5.43E-01 | -0.0095 | 0.0086 | 2.69E-01 | -0.0073 | 0.0092 | 4.28E-01 | -0.0026 | 0.0087 | 7.65E-01 |
| | DRB1*0801 | 1.82% | 0.0109 | 0.0193 | 5.72E-01 | -0.0415 | 0.0184 | 2.44E-02 | 0.0163 | 0.0193 | 4.00E-01 | -0.0371 | 0.0185 | 4.44E-02 |
| | DRB1*0901 | 1.28% | 0.0004 | 0.0222 | 9.84E-01 | -0.0134 | 0.0214 | 5.31E-01 | -0.0015 | 0.0222 | 9.48E-01 | -0.0042 | 0.0214 | 8.43E-01 |
| | DRB1*1001 | 0.59% | 0.0318 | 0.0329 | 3.34E-01 | -0.0420 | 0.0308 | 1.74E-01 | 0.0318 | 0.0323 | 3.25E-01 | -0.0389 | 0.0303 | 1.99E-01 |
| | DRB1*1101 | 3.50% | 0.0071 | 0.0145 | 6.24E-01 | -0.0185 | 0.0139 | 1.84E-01 | 0.0076 | 0.0147 | 6.04E-01 | -0.0110 | 0.0140 | 4.34E-01 |
| | DRB1*1104 | 1.16% | 0.0440 | 0.0236 | 6.17E-02 | -0.0727 | 0.0225 | 1.21E-03 | 0.0464 | 0.0237 | 5.05E-02 | -0.0638 | 0.0226 | 4.73E-03 |
| | DRB1*1201 | 1.45% | 0.0265 | 0.0213 | 2.13E-01 | -0.0303 | 0.0204 | 1.37E-01 | 0.0267 | 0.0214 | 2.11E-01 | -0.0261 | 0.0204 | 2.01E-01 |
| | DRB1*1301 | 5.05% | 0.0078 | 0.0126 | 5.34E-01 | -0.0077 | 0.0120 | 5.22E-01 | 0.0096 | 0.0127 | 4.51E-01 | -0.0017 | 0.0121 | 8.92E-01 |
| | DRB1*1302 | 3.82% | 0.0062 | 0.0140 | 6.60E-01 | -0.0190 | 0.0134 | 1.56E-01 | 0.0061 | 0.0143 | 6.71E-01 | -0.0093 | 0.0137 | 4.98E-01 |
| | DRB1*1303 | 0.84% | -0.0423 | 0.0272 | 1.20E-01 | 0.0287 | 0.0263 | 2.76E-01 | -0.0378 | 0.0269 | 1.59E-01 | 0.0307 | 0.0260 | 2.39E-01 |

| | | | | | | | | | | | | | | |
|---|---|---|---|---|---|---|---|---|---|---|---|---|---|---|
| | DRB1*1401 | 1.92% | -0.0172 | 0.0188 | 3.60E-01 | 0.0047 | 0.0179 | 7.93E-01 | -0.0128 | 0.0190 | 5.01E-01 | 0.0116 | 0.0181 | 5.22E-01 |
| | DRB1*1501 | 14.62% | 0.0098 | 0.0090 | 2.76E-01 | -0.0081 | 0.0086 | 3.50E-01 | 0.0097 | 0.0090 | 2.80E-01 | -0.0062 | 0.0086 | 4.70E-01 |
| | DRB1*1601 | 0.55% | -0.0178 | 0.0337 | 5.97E-01 | -0.0011 | 0.0321 | 9.74E-01 | -0.0085 | 0.0336 | 8.01E-01 | 0.0005 | 0.0321 | 9.87E-01 |
| | DRB1*Other | 2.58% | -0.0179 | 0.0143 | 2.11E-01 | 0.0054 | 0.0138 | 6.96E-01 | -0.0179 | 0.0146 | 2.20E-01 | 0.0141 | 0.0139 | 3.11E-01 |
| DRB3 | rs9366778 | NA | NA | NA | NA | NA | NA | NA | 0.0031 | 0.0053 | 5.64E-01 | -0.0199 | 0.0051 | 8.67E-05 |
| | rs6911024 | NA | NA | NA | NA | NA | NA | NA | -0.0415 | 0.0086 | 1.54E-06 | 0.0130 | 0.0083 | 1.17E-01 |
| | DRB3*0101 | 16.79% | -0.0003 | 0.0066 | 9.64E-01 | 0.0080 | 0.0063 | 2.06E-01 | -0.0013 | 0.0067 | 8.45E-01 | 0.0031 | 0.0065 | 6.36E-01 |
| | DRB3*0202 | 13.38% | 0.0116 | 0.0072 | 1.07E-01 | -0.0062 | 0.0069 | 3.72E-01 | 0.0117 | 0.0072 | 1.04E-01 | -0.0049 | 0.0069 | 4.82E-01 |
| | DRB3*0301 | 3.82% | 0.0080 | 0.0127 | 5.27E-01 | -0.0103 | 0.0122 | 3.97E-01 | 0.0080 | 0.0127 | 5.28E-01 | -0.0063 | 0.0122 | 6.07E-01 |
| DRB4 | rs9366778 | NA | NA | NA | NA | NA | NA | NA | 0.0067 | 0.0053 | 2.02E-01 | -0.0208 | 0.0051 | 3.85E-05 |
| | rs6911024 | NA | NA | NA | NA | NA | NA | NA | -0.0430 | 0.0088 | 9.62E-07 | 0.0133 | 0.0084 | 1.15E-01 |
| | DRB4*0101 | 8.34% | -0.0088 | 0.0089 | 3.21E-01 | -0.0082 | 0.0085 | 3.33E-01 | -0.0142 | 0.0091 | 1.16E-01 | -0.0009 | 0.0087 | 9.20E-01 |
| | DRB4*0103 | 25.94% | 0.0015 | 0.0056 | 7.94E-01 | -0.0013 | 0.0054 | 8.05E-01 | -0.0021 | 0.0056 | 7.04E-01 | 0.0013 | 0.0054 | 8.05E-01 |
| DRB5 | rs9366778 | NA | NA | NA | NA | NA | NA | NA | 0.0061 | 0.0052 | 2.42E-01 | -0.0219 | 0.0050 | 1.07E-05 |
| | rs6911024 | NA | NA | NA | NA | NA | NA | NA | -0.0412 | 0.0086 | 1.83E-06 | 0.0138 | 0.0083 | 9.71E-02 |
| | DRB5*0101 | 14.57% | 0.0110 | 0.0068 | 1.08E-01 | 0.0016 | 0.0065 | 8.07E-01 | 0.0097 | 0.0069 | 1.62E-01 | -0.0025 | 0.0066 | 7.11E-01 |
| | DRB5*0202 | 0.71% | -0.0272 | 0.0289 | 3.46E-01 | 0.0065 | 0.0274 | 8.13E-01 | -0.0162 | 0.0288 | 5.75E-01 | 0.0015 | 0.0273 | 9.56E-01 |

The “m” and “f” prefix of variable name indicate the maternal and fetal effects of the corresponding allele, respectively; SE: standard error; [1] The effect alleles for rs9366778 is A and the effect allele for rs6911024 is C.

Supplementary Figures S1-5

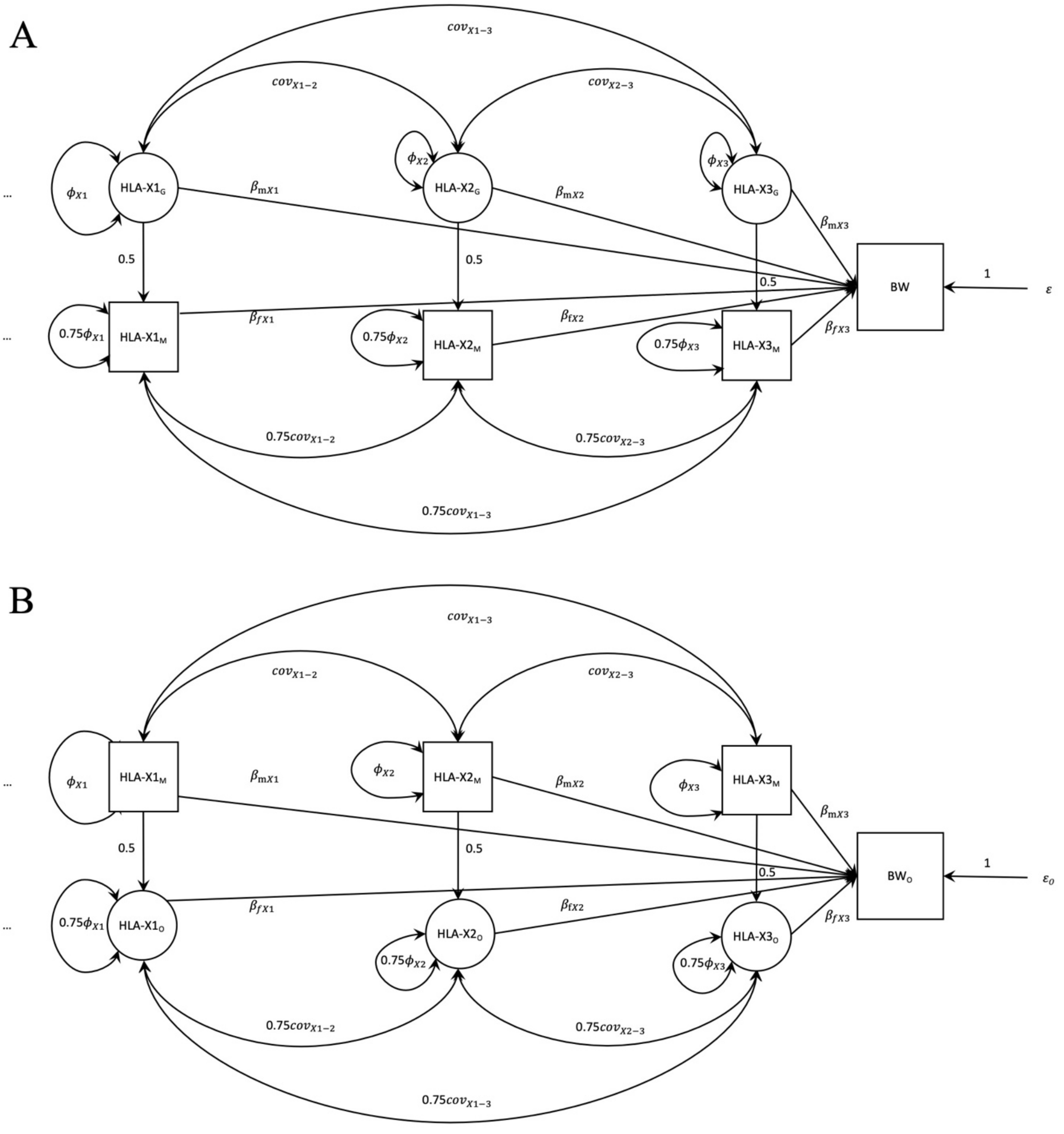


Figure S1 A two component structural equation model (SEM) used for the analysis of multi-allelic HLA markers and birth weight in which some genotyped individuals only report their own birth weight and some genotyped individuals only report their offspring's birth weight.

Panel A illustrates the component which models genotyped individuals who only report their own birth weight; Panel B illustrates the component which models individuals who only report the birth weight of their first-born. Latent variables are represented by circles, observed variables by square boxes. Causal paths are indicated by unidirectional arrows, bidirectional arrows indicate covariances. In order to illustrate the model we use the fictional example of the *HLA-X* gene. We assume there are only four alleles in the *HLA-X* gene (here *HLA-X0*, *HLA-X1*, *HLA-X2*, *HLA-X3*), although the model is generalizable to an arbitrary number of alleles (represented by the ellipsis "…" on the left-hand side of the path model). The *HLA-X0* allele is assumed to be the most common allele and is modelled as the "baseline" (so all effects are modelled as additive displacements from the baseline homozygous *HLA-X0* genotype), and hence is not shown in the diagram. The *HLA-X1*, *HLA-X2*, and *HLA-X3* variables represent the number of copies of each of these alleles that an individual carries. The "G", "M", and "O" subscripts index genotypes in the grandmaternal (latent), maternal (observed), and offspring (latent) generation respectively. The "X1", "X2", and "X3" subscripts index the corresponding alleles. The "m" and "f" subscripts of the path coefficients represent maternal and fetal genetic effects, respectively. The observed variables, displayed in squares, in the analysis are 1) the self-reported birth weight of the individual (BW) in the UKB, 2) the self-reported birth weight of their first offspring in the case of women in the UK Biobank ($BW_O$), and 3) the number of copies of the *HLA-X1* ($HLA\text{-}X1_M$), *HLA-X2* ($HLA\text{-}X2_M$), and *HLA-X3* ($HLA\text{-}X3_M$) alleles. The latent variables, displayed in circles, in the analysis are 1) the number of copies of the relevant allele in the genotyped individual's mother (i.e. $HLA\text{-}X1_G$, $HLA\text{-}X2_G$, and $HLA\text{-}X3_G$) and 2) the number of copies of the relevant allele in the genotyped individual's offspring ($HLA\text{-}X1_O$, $HLA\text{-}X2_O$, and $HLA\text{-}X3_O$). The variance of the allele counts in the grandmaternal ($HLA\text{-}X1_G$, $HLA\text{-}X2_G$, $HLA\text{-}X3_G$), maternal ($HLA\text{-}X1_M$, $HLA\text{-}X2_M$, $HLA\text{-}X3_M$) and offspring ($HLA\text{-}X1_O$, $HLA\text{-}X2_O$, $HLA\text{-}X3_O$) generations are estimated as $\Phi_{X1}$, $\Phi_{X2}$ and $\Phi_{X3}$, respectively, and assumed not to change across generations. The covariances between the different allele counts are also estimated (e.g. $cov_{X1\text{-}2}$, $cov_{X1\text{-}3}$, $cov_{X2\text{-}3}$) and assumed not to vary across generations. The $\beta_{mX1}$, $\beta_{fX1}$, $\beta_{mX2}$, $\beta_{fX2}$, $\beta_{mX3}$ and $\beta_{fX3}$ are path coefficients that quantify the maternal and fetal genetic effects of each allele on birth weight, respectively. The residual error terms for the birth weight of the individual and their offspring are represented by $\varepsilon$ and $\varepsilon_o$ respectively, and the variance of both of these terms is estimated in the SEM.

**UKB BW phenotype/sample cleaning**

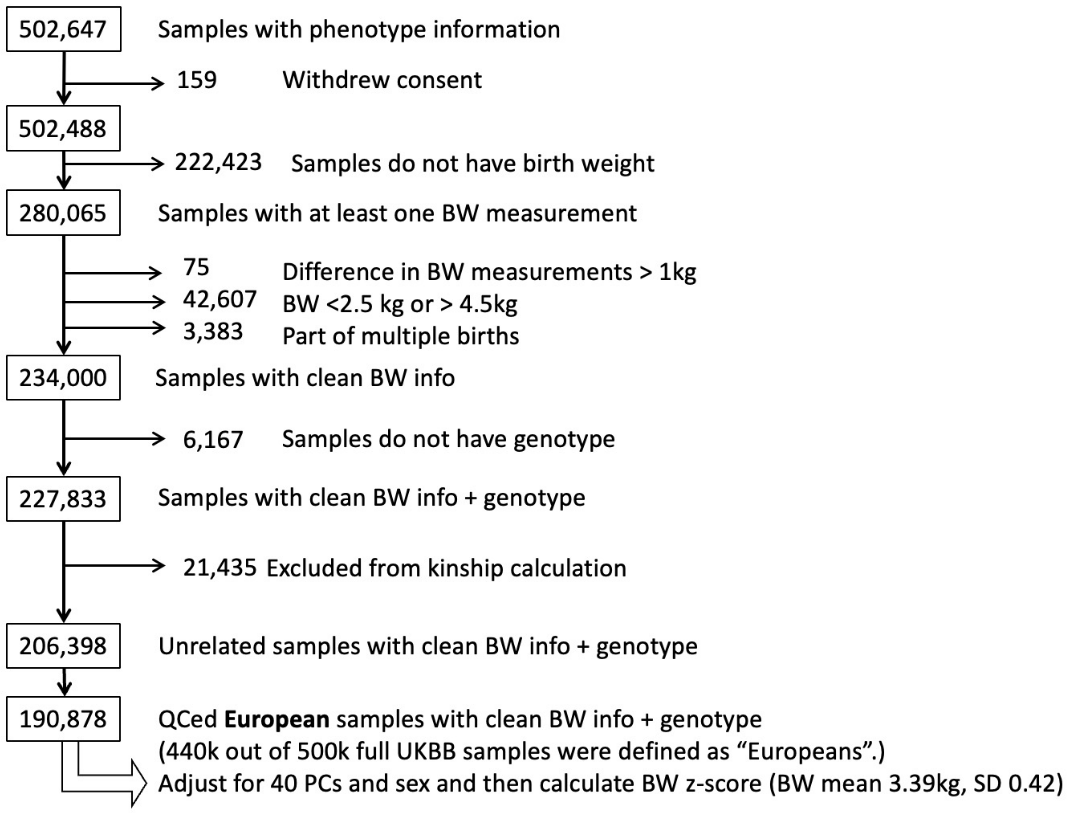


Figure S2 Flow chart of data cleaning of birth weight phenotype from UK Biobank

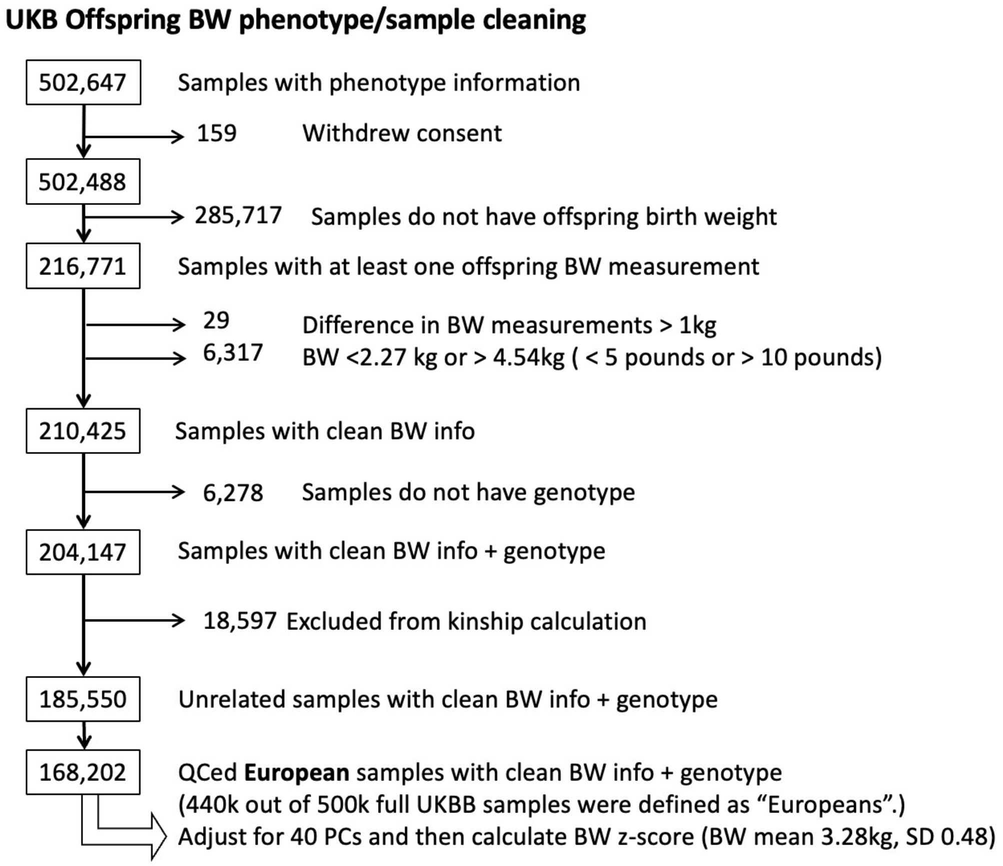


Figure S3 Flow chart of data cleaning of offspring birth weight phenotype from UK Biobank

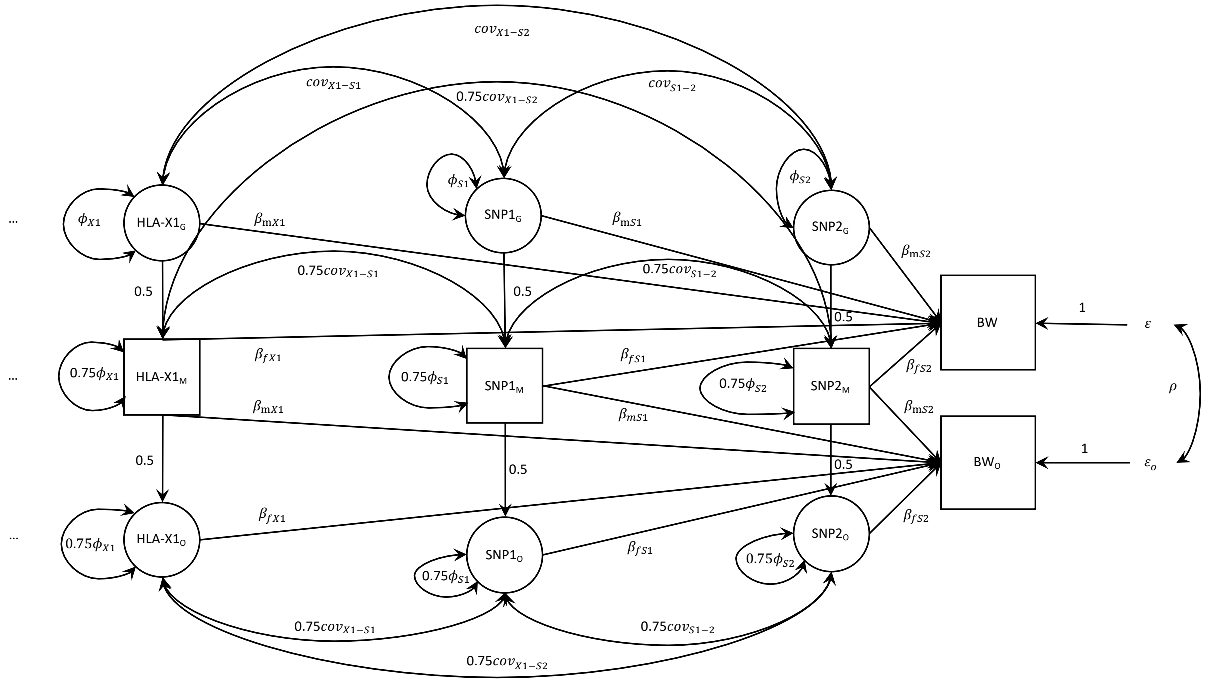


Figure S4 Structural equation model (SEM) used for the conditional analysis of multi-allelic HLA markers and birth weight conditioning on GWAS-associated SNPs.

Latent variables are represented by circles, observed variables by square boxes. Causal paths are indicated by unidirectional arrows, bidirectional arrows represent covariances. In order to illustrate the model we use the fictional example of the *HLA-X* gene with an arbitrary number of alleles (*HLA-X0*, *HLA-X1, HLA-X2*, *HLA-X3...*), and SNP1 and SNP2 (representing the SNPs rs9366778 and rs6911024 in the empirical analysis). The diagram only shows one classical HLA allele *HLA-X1* for illustrative purposes, although the model is generalizable to an arbitrary number of alleles (represented by the ellipsis "…" on the left-hand side of the path model, e.g., *HLA-X2*, *HLA-X3...*). The *HLA-X0* allele is modelled as the "baseline" allele (so all effects are modelled as displacements from this baseline genotype), and hence is not shown in the diagram. The *HLA-X1* variable represents the number of copies of the alleles that each individual carries. The SNP1 and SNP2 variables represent the number of copies of the effect allele at each SNP that each individual carries. The "G", "M", and "O" subscripts index genotypes in the grandmaternal (latent), maternal (observed), and offspring (latent) generation respectively. The "X1", "S1", and "S2" subscripts refer to the *HLA-X1*, SNP1 and SNP2 variables respectively. The "m" and "f" subscripts of the path coefficients refer to maternal and fetal genetic effects, respectively. The five observed variables, displayed in squares, in the analysis are 1) the self-reported birth weight of the individual (BW) in the UKB, 2) the self-reported birth weight of their first offspring (in the case of women only) in the UK Biobank ($BW_O$), and 3) the number of copies of the *HLA-X1* ($HLA\text{-}X1_M$) allele, and genotypes of SNP1 ($SNP1_M$), and SNP2 ($SNP2_M$). The latent variables, displayed in circles, in the analysis are 1) the number of copies of the relevant allele carried by the individual's mother (i.e. $HLA\text{-}X1_G$, $SNP1_G$, $SNP2_G$) and 2) the number of copies of the relevant allele carried by the individual's offspring ($HLA\text{-}X1_O$, $SNP1_O$, $SNP2_O$). The variance of the allele counts in the grandmaternal ($HLA\text{-}X1_G$, $SNP1_G$, $SNP2_G$), maternal ($HLA\text{-}X1_M$, $SNP1_M$, $SNP2_M$) and offspring ($HLA\text{-}X1_O$, $SNP1_O$, $SNP2_O$) generations are estimated as $\Phi_{X1}$, $\Phi_{S1}$ and $\Phi_{S2}$, respectively, and assumed not to change across generations (i.e., variance ($HLA\text{-}X_G$) = Φ, variance ($HLA\text{-}X_M$) = 0.75Φ + 0.25Φ = Φ, and variance ($HLA\text{-}X_O$) = 0.75Φ + 0.25Φ = Φ). The covariances across the *HLA-X1* allele, SNP1, and SNP2 are also estimated (e.g. $cov_{X1\text{-}S1}$, $cov_{X1\text{-}S2}$, $cov_{S1\text{-}2}$) and assumed not to vary across generations (e.g., covariance ($HLA\text{-}X1_G$, $SNP1_G$) = $cov_{X1\text{-}S1}$, covariance ($HLA\text{-}X1_M$, $SNP1_M$) = 0.75 $cov_{X1\text{-}S1}$+ 0.25 $cov_{X1\text{-}S1}$= $cov_{X1\text{-}S1}$, and covariance ($HLA\text{-}X1_O$, $SNP1_O$) = 0.75 $cov_{X1\text{-}S1}$+ 0.25 $cov_{X1\text{-}S1}$= $cov_{X1\text{-}S1}$).. The $\beta_{mX1}$, $\beta_{fX1}$, $\beta_{mS1}$, $\beta_{fS1}$, $\beta_{mS2}$ and $\beta_{fS2}$ are path coefficients that quantify the maternal and fetal genetic effects of the *X1* allele and each SNP on birth weight. The residual error terms for the birth weight of the individual and their offspring are represented by $\varepsilon$ and $\varepsilon_o$ respectively, and the variance of both of these terms is estimated in the SEM. The estimated covariance between residual genetic and environmental sources of variation on birth weight is represented by $\rho$.

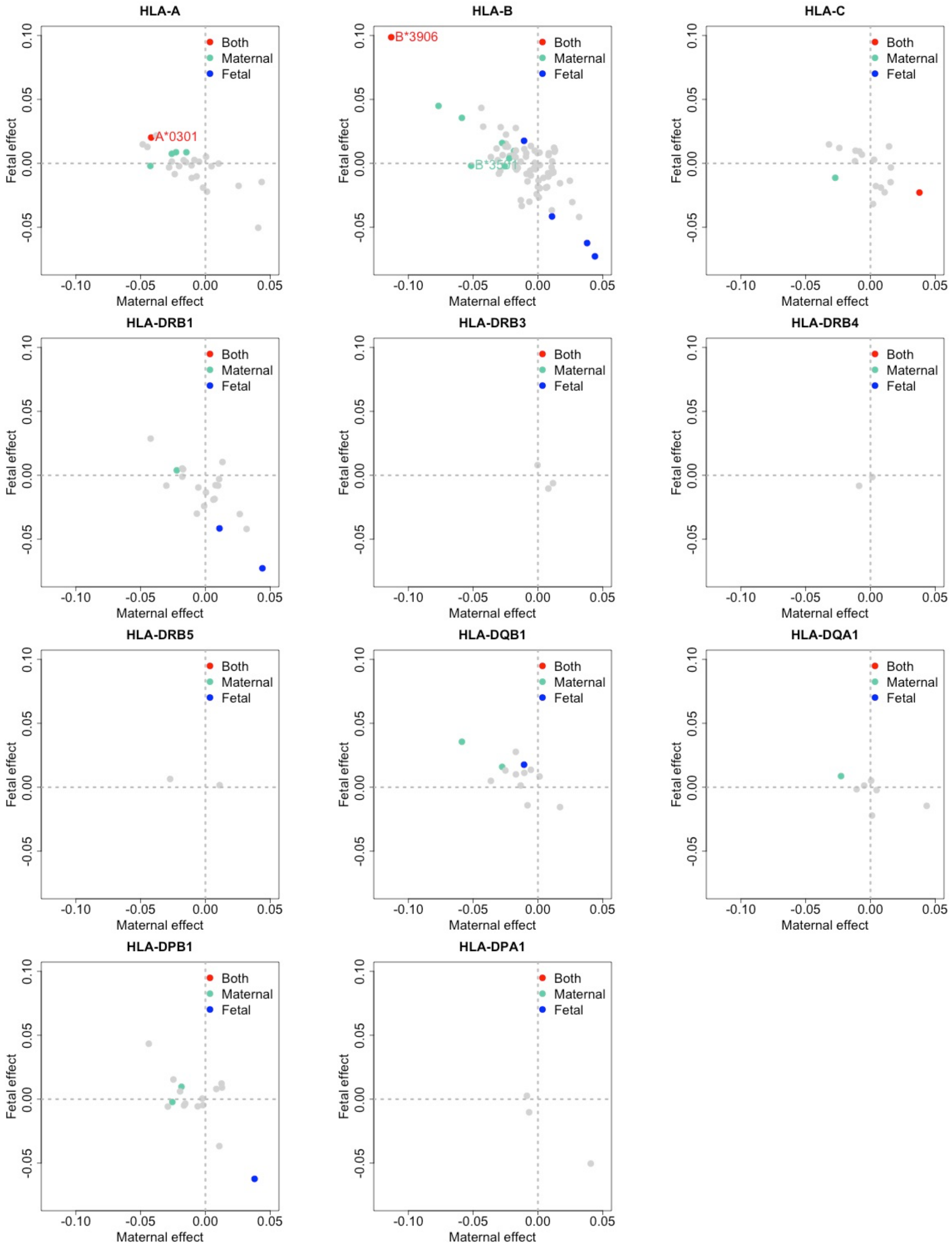


Figure S5 HLA associations with birth weight in UK Biobank at each locus

Figure S5 Maternal and fetal effect estimates for classical HLA alleles and birth weight in the UK Biobank at each locus.

Fetal and maternal effect sizes were estimated using the structural equation model. The colour of each dot represents whether maternal (green) and/or fetal associations (blue) passed nominal significance ($P < 0.05$). Red dots indicate alleles where tests for both maternal and fetal effects reached the threshold. Light grey dots represent alleles which did not $P < 0.05$. Alleles with $P < 0.001$ are explicitly labelled. The observed negative correlation between maternal and fetal effects is at least partially a consequence of the negative correlation between these parameters in the SEM.

Supplementary Note

Example R script

```
library("OpenMx")
#Submodel1 for data have both individuals' own and offspring birth weight
#data_sub is a data frame contains individuals' own and offspring birth weight (bm and bo)
and HLA alleles
#"bm" "bo" A_101" "A_102" "A_103" "A_202" "A_203" ...
# 2.5  2.5   1      1       0       0       0      ...
############################RAM model ##matrix adhesion
manifests <- names(data_sub) ##names of observed variables
modelData <- data_sub ## data set
nvar <- ncol(modelData) ##number of observed variables
varnames <- colnames(modelData) #
nvarall <- nvar + 6*(nvar-2) +2 #number of variables

latents<-c()
for (i in  1:(nvar-2)){
  latents[(6*(i-1)+1):(6*i)]=
    c(paste0("gvar",i),
      paste0( varnames[i+2],"_g"),
      paste0("mvar",i),
      paste0(varnames[i+2],"_m"),
      paste0("ovar",i),
      paste0(varnames[i+2],"_o"))
}
latents<-c(latents,"e","eo")   ##names of latent variables


allVariables <- c(varnames,latents) ##all variables

##A matrix

###Construct A matrix (details of  A , S , F , M matrices ?mxRAMObjective())
#The 'A' argument refers to the A or asymmetric matrix in the RAM approach. This matrix
consists of all of the asymmetric paths (one-headed arrows) in the model. A free parameter in
any row and column describes a regression of the variable represented by that row regressed
on the variable represented in that column.
A1 <- mxMatrix("Zero", nrow=nvarall,ncol=nvar,labels=NA, byrow=TRUE, name="A1")
A_beta <- mxMatrix('Zero', nvar,0)
##A2-A3
for (i in 2:(nvar-1)){
  assign(paste0("A",i),mxMatrix("Full", nrow=nvar,ncol=6,
                              labels =
c(c(NA,paste0("m",varnames[i+1]),NA,paste0("f",varnames[i+1]),NA,NA, #"m" indicates
maternal effect
                                     NA,NA,NA,
paste0("m",varnames[i+1]),NA,paste0("f",varnames[i+1])), #"f" indicates fetal effect
                                   rep(NA, 6*(nvar-2))),
```

```
                          values = c(c(rep(0,i*6+3)),
                                   1,c(rep(0,(6*(nvar-2)-(i-2)*6-4)))),
                          free = c(c(F,T,F,T,F,F,
                                   F,F,F,T,F,T),
                                 c(rep(F,6*(nvar-2)))),
                          byrow=TRUE,name="A2"))
  A_beta<-omxCbind(A_beta,get(paste0("A",i)), name="A_beta")
}


A_mid <-  mxMatrix("Full", nrow=6,ncol=6,
                labels = NA,
                values = c(c(rep(0,6)),
                         1,
                         c(rep(0,12)),
                         0.5,sqrt(0.75),
                         c(rep(0,12)),
                         0.5,sqrt(0.75),0),
                free = F,
                name="Amid",
                byrow=TRUE)
A_diag <-  mxMatrix("Full", nrow=6,ncol=6,
                labels = NA,
                values = c(c(rep(0,6)),
                         1,
                         c(rep(0,12)),
                         0.5,sqrt(0.75),
                         c(rep(0,12)),
                         0.5,sqrt(0.75),0),
                free = F,
                name="Ad",
                byrow=TRUE)
for (i in 1:(nvar-3)){

  A_zeroh <- mxMatrix("Zero", nrow=6,ncol=6*i,labels=NA, byrow=TRUE, name="Azh")
  A_zerov <- mxMatrix("Zero", nrow=6*i,ncol=6,labels=NA, byrow=TRUE, name="Azv")

  A_mid <- omxRbind(omxCbind(A_mid, A_zerov),
                omxCbind(A_zeroh,A_diag),name= "A_mid")
}


A_error <- mxMatrix("Full", nrow=nvarall,ncol=2,labels=NA, free = F,
                values = c(1,0,
                         0,1,
                         rep(0,2*nvarall-4)), byrow=TRUE, name="Ae")
A_bottom <- mxMatrix("Zero", nrow=2,ncol=6*(nvar-2),labels=NA, byrow=TRUE,
name="Ab")
```

```
Am <- omxCbind(A1,omxRbind(A_beta,
                   A_mid,A_bottom),A_error,name="A",
          dimnames = list(allVariables,allVariables))
```

##S matrix

#The 'S' argument refers to the S or symmetric matrix in the RAM approach, and as such must be square. This matrix consists of all of the symmetric paths (two-headed arrows) in the model. A free parameter in any row and column describes a covariance between the variable represented by that row and the variable represented by that column. Variances are covariances between any variable at itself, which occur on the diagonal of the specified matrix.

```
S1 <-  mxMatrix("Zero", nrow=nvarall,ncol=nvar,labels=NA, byrow=TRUE, name="S1")
S_top <- mxMatrix("Zero", nrow=nvar,ncol=6*(nvar-2),labels=NA, byrow=TRUE,
name="St")


S_mid <- mxMatrix("Diag", nrow=6,ncol=6,
            labels=c(paste0("var_",varnames[3]),NA, ##"var_" indicates the variance term
                  paste0("var_",varnames[3]),NA,
                  paste0("var_",varnames[3]),NA),
            values = c(0.2, 0,0.2, 0,0.2, 0),
            free= c(T,F,T,F,T,F),
            byrow=TRUE, name="Sd")
for (i in 2:(nvar-2)){
  S_diag <- mxMatrix("Diag", nrow=6,ncol=6,
              labels=c(paste0("var_",varnames[i+2]),NA,
                    paste0("var_",varnames[i+2]),NA,
                    paste0("var_",varnames[i+2]),NA),
              values = c(0.2, 0, 0.2, 0, 0.2, 0),
              free= c(T,F,T,F,T,F),
              byrow=TRUE, name="Sd")
  S_covv <-mxMatrix("Zero", nrow=0,ncol=6,labels=NA, byrow=TRUE, name="Sch")
  S_covh <-mxMatrix("Zero", nrow=6,ncol=0,labels=NA, byrow=TRUE, name="Sch")
  for (j in 2:i){
    S_covh <- omxCbind( S_covh , mxMatrix("Diag", nrow=6,ncol=6,

labels=c(rep(c(paste0("cov_",varnames[j+1],"_",varnames[i+2]),NA),3)), ##"cor_" indicates
the covariance term
                             values = 0,
                             free= c(T,F,T,F,T,F),
                             byrow=TRUE, name="S_ch"))
    S_covv <- omxRbind( S_covv , mxMatrix("Diag", nrow=6,ncol=6,

labels=c(rep(c(paste0("cov_",varnames[j+1],"_",varnames[i+2]),NA),3)),
                             values = 0,
```

```
                                free= c(T,F,T,F,T,F),
                                byrow=TRUE, name="S_cv"))
  }


  S_mid <-  omxRbind(omxCbind(S_mid, S_covv),
              omxCbind(S_covh, S_diag))
}


S_error <- mxMatrix("Full", nrow=nvarall,ncol=2,
              values = c(rep(0,2*nvarall-4),
                    1,0.2,
                    0.2,1),
              free= c(rep(F,2*nvarall-4),T,T,T,T),
              labels = c(rep(NA,2*nvarall-4),"phi1", "rho", "rho", "phi2"),
              ##"phi1" and "phi2" are error terms of "bm" and "bo"
              ##"rho" is the convariance between "phi1" and "phi2"
              byrow=TRUE, name="Se")
S_bottom <- mxMatrix("Zero", nrow=2,ncol=6*(nvar-2),labels=NA, byrow=TRUE,
name="Sb")
Sm<- omxCbind(S1,
          omxRbind(S_top,S_mid,S_bottom),
          S_error,
          name="S", dimnames = list(allVariables,allVariables))
```

##F matrix
#The 'F' argument refers to the F or filter matrix in the RAM approach. If no latent variables are included in the model (i.e., the A and S matrices are of both of the same dimension as the data matrix), then the 'F' should refer to an identity matrix. If latent variables are included (i.e., the A and S matrices are not of the same dimension as the data matrix), then the 'F' argument should consist of a horizontal adhesion of an identity matrix and a matrix of zeros.

```
F1 <- mxMatrix("Iden", nrow=nvar,ncol=nvar,labels = NA,byrow=TRUE, name="F1")
F2 <- mxMatrix("Zero", nrow=nvar,ncol=nvarall-nvar, labels =NA,byrow=TRUE,
name="F2")
Fm <-  omxCbind(F1,F2,name="F", dimnames = list(manifests,allVariables))
```

##M matrix
#The 'M' argument refers to the M or means matrix in the RAM approach. It is a 1 x n matrix, where n is the number of manifest variables + the number of latent variables. The M matrix must be specified if either the mxData type is “cov” or “cor” and a means vector is provided, or if the mxData type is “raw”. Otherwise the M matrix is ignored.

```
Mm <- mxMatrix("Full", nrow=1,ncol=nvarall,dimnames=list(NULL,allVariables),
          values= 0,
          free=c(rep(T,nvar),rep(F,nvarall-nvar)),
          labels=c(paste0("mean_",varnames),rep(NA,nvarall-nvar)) ##"mean_" indicates the
means of each observed variables
          , byrow=TRUE, name="M")
```

```
fitFunction <- mxFitFunctionML()
expFunction <- mxExpectationRAM(A ="A",S="S",F = "F",M ="M",
                          dimnames= allVariables)
#Model1
model_1 <- mxModel("SEM_BW_HLA",
             type= "RAM",
             Am,Sm,Fm,Mm,
             mxData(modelData, type = "raw"),
             expFunction,
             fitFunction )
##################################################################3
#Submodel2 for data only has individuals' own birth weight
#data_sub2 contain individuals' own birth weight (bm) and HLA alleles
#"bm" "A_101" "A_102" "A_103" "A_202" "A_203" ...
# 0.1  1     1       0      0      0     ...

manifests2 <- names(data_sub2) ##annotations same as above in Submodel1
modelData2 <- data_sub2
nvar2 <- ncol(modelData2)
varnames2 <- colnames(modelData2)
nvarall2 <- nvar2 + 4*(nvar2-1) +1

latents2<-c()
for (i in  1:(nvar2-1)){
  latents2[(4*(i-1)+1):(4*i)]=
    c(paste0("gvar",varnames2[i+1]),
      paste0( "A",varnames2[i+1],"_g"),
      paste0("mvar",varnames2[i+1]),
      paste0("A",varnames2[i+1]))
}
latents2<-c(latents2,"e")


allVariables2 <- c(varnames2,latents2)



##A2-A3
A1_2 <- mxMatrix("Zero", nrow=nvarall2,ncol=nvar2,labels=NA, byrow=TRUE,
name="A1_2")
A_beta_2 <- mxMatrix('Zero', nvar2,0)
##A2-A3
for (i in 2:(nvar2)){
  assign(paste0("A",i,"_2"),mxMatrix("Full", nrow=nvar2,ncol=4,
                          labels =
c(c(NA,paste0("m",varnames2[i]),NA,paste0("f",varnames2[i])),

                                 rep(NA, 4*(nvar2-1))),
                         values = c(c(rep(0,(i-1)*4+3)),
                                 1,c(rep(0,(4*(nvar2-1)-(i-2)*4-4)))),
                         free = c(c(F,T,F,T),
```

```
                                   c(rep(F,4*(nvar2-1)))),
                              byrow=TRUE,name="A2_2"))
  A_beta_2<-omxCbind(A_beta_2,get(paste0("A",i,"_2")), name="A_beta_2")
}


A_mid_2 <-  mxMatrix("Full", nrow=4,ncol=4,
                labels = NA,
                values = c(c(rep(0,4)),
                        1,
                        c(rep(0,8)),
                        0.5,sqrt(0.75),
                        0),
                free = F,
                name="Amid_2",
                byrow=TRUE)
A_diag_2 <-  mxMatrix("Full", nrow=4,ncol=4,
                 labels = NA,
                 values = c(c(rep(0,4)),
                         1,
                         c(rep(0,8)),
                         0.5,sqrt(0.75),
                         0),
                 free = F,
                 name="Ad_2",
                 byrow=TRUE)
for (i in 1:(nvar2-2)){

  A_zeroh_2 <- mxMatrix("Zero", nrow=4,ncol=4*i,labels=NA, byrow=TRUE,
name="Azh_2")
  A_zerov_2 <- mxMatrix("Zero", nrow=4*i,ncol=4,labels=NA, byrow=TRUE,
name="Azv_2")

  A_mid_2 <- omxRbind(omxCbind(A_mid_2, A_zerov_2),
                 omxCbind(A_zeroh_2,A_diag_2),name= "A_mid_2")
}


A_error_2 <- mxMatrix("Full", nrow=nvarall2,ncol=1,labels=NA, free = F,
                 values = c(1,
                         rep(0,nvarall2-1)), byrow=TRUE, name="Ae_2")
A_bottom_2 <- mxMatrix("Zero", nrow=1,ncol=4*(nvar2-1),labels=NA, byrow=TRUE,
name="Ab_2")




Am_2 <- omxCbind(A1_2,omxRbind(A_beta_2,
                        A_mid_2,A_bottom_2),A_error_2,name="A",
             dimnames = list(allVariables2,allVariables2))
```

##S matrix

```
S1_2 <-  mxMatrix("Zero", nrow=nvarall2,ncol=nvar2,labels=NA, byrow=TRUE,
name="S1_2")
S_top_2 <- mxMatrix("Zero", nrow=nvar2,ncol=4*(nvar2-1),labels=NA, byrow=TRUE,
name="St_2")


S_mid_2 <- mxMatrix("Diag", nrow=4,ncol=4,
              labels=c(paste0("var_",varnames2[2]),NA,
                    paste0("var_",varnames2[2]),NA
              ),
              values = c(0.2, 0, 0.2, 0),
              free= c(T,F,T,F),
              byrow=TRUE, name="Smid_2")
for (i in 2:(nvar2-1)){
  S_diag_2 <- mxMatrix("Diag", nrow=4,ncol=4,
                labels=c(paste0("var_",varnames2[i+1]),NA,
                      paste0("var_",varnames2[i+1]),NA),
                values = c(0.2, 0, 0.2, 0),
                free= c(T,F,T,F),
                byrow=TRUE, name="Sd_2")
  S_covv_2 <-mxMatrix("Zero", nrow=0,ncol=4,labels=NA, byrow=TRUE, name="Sch_2")
  S_covh_2 <-mxMatrix("Zero", nrow=4,ncol=0,labels=NA, byrow=TRUE, name="Sch_2")
  for (j in 2:i){
    S_covh_2 <- omxCbind( S_covh_2 , mxMatrix("Diag", nrow=4,ncol=4,

labels=c(rep(c(paste0("cov_",varnames2[j],"_",varnames2[i+1]),NA),2)),
                                values = 0,
                                free= c(T,F,T,F),
                                byrow=TRUE, name="S_ch_2"))
    S_covv_2 <- omxRbind( S_covv_2 , mxMatrix("Diag", nrow=4,ncol=4,

labels=c(rep(c(paste0("cov_",varnames2[j],"_",varnames2[i+1]),NA),2)),
                                values = 0,
                                free= c(T,F,T,F),
                                byrow=TRUE, name="S_cv_2"))
  }


  S_mid_2 <-  omxRbind(omxCbind(S_mid_2, S_covv_2),
                omxCbind(S_covh_2, S_diag_2))
}



S_error_2 <- mxMatrix("Full", nrow=nvarall2,ncol=1,
```

```
                  values = c(rep(0,nvarall2-1),
                          1),
                  free= c(rep(F,nvarall2-1),T),
                  labels = c(rep(NA,nvarall2-1),"phi1"),
                  byrow=TRUE, name="Se_2")
S_bottom_2 <- mxMatrix("Zero", nrow=1,ncol=4*(nvar2-1),labels=NA, byrow=TRUE,
name="Sb_2")
Sm_2<- omxCbind(S1_2,
          omxRbind(S_top_2,S_mid_2,S_bottom_2),
          S_error_2,
          name="S", dimnames = list(allVariables2,allVariables2))

##F matrix
F1_2 <- mxMatrix("Iden", nrow=nvar2,ncol=nvar2,labels = NA,byrow=TRUE,
name="F1_2")
F2_2 <- mxMatrix("Zero", nrow=nvar2,ncol=nvarall2-nvar2, labels =NA,byrow=TRUE,
name="F2_2")
Fm_2 <-  omxCbind(F1_2,F2_2,name="F", dimnames = list(manifests2,allVariables2))

##M matrix
Mm_2 <- mxMatrix("Full", nrow=1,ncol=nvarall2,dimnames=list(NULL,allVariables2),
          values= 0,
          free=c(rep(T,nvar2),rep(F,nvarall2-nvar2)),
          labels=c(paste0("mean_",varnames2),rep(NA,nvarall2-nvar2))
          , byrow=TRUE, name="M")
fitFunction_2 <- mxFitFunctionML()
expFunction_2 <- mxExpectationRAM(A="A", S="S", F="F", M = "M",
                    dimnames= allVariables2)

model_2 <- mxModel("SEM_BW_HLA_2",
           type= "RAM",
           Am_2,Sm_2,Fm_2, Mm_2,
           mxData(modelData2, type = "raw"),
           expFunction_2,
           fitFunction_2 )


#########################################################################
#Submodel3 for data only has individuals' offspring birth weight
#data_sub3 contain individuals' offspring birth weight (bo) and HLA alleles
#"bo" "A_101" "A_102" "A_103" "A_202" "A_203" ...
# 0.1  1    1     0     0     0    ...
# ...

manifests3 <- names(data_sub3)
modelData3 <- data_sub3
nvar3 <- ncol(modelData3)
varnames3 <- colnames(modelData3)
nvarall3 <- nvar3 + 4*(nvar3-1) +1
```

```
latents3<-c()
for (i in  1:(nvar3-1)){
  latents3[(4*(i-1)+1):(4*i)]=
    c(
      paste0("mvar",varnames3[i+1]),
      paste0("A",varnames3[i+1]),
      paste0("ovar",varnames3[i+1]),
      paste0("A",varnames3[i+1],"_o"))
}
latents3<-c(latents3,"eo")


allVariables3 <- c(varnames3,latents3)


##A2-A3
#######
A1_3 <- mxMatrix("Zero", nrow=nvarall3,ncol=nvar3,labels=NA, byrow=TRUE,
name="A1_3")
A_beta_3 <- mxMatrix('Zero', nvar3,0)
##A2-A3
for (i in 2:(nvar3)){
  assign(paste0("A",i,"_3"),mxMatrix("Full", nrow=nvar3,ncol=4,
                                    labels =
c(c(NA,paste0("m",varnames3[i]),NA,paste0("f",varnames3[i])),

                                               rep(NA, 4*(nvar3-1))),
                                    values = c(c(rep(0,(i-1)*4+1)),
                                               1,c(rep(0,(4*(nvar3-1)-(i-2)*4-2)))),
                                    free = c(c(F,T,F,T),
                                             c(rep(F,4*(nvar3-1)))),
                                    byrow=TRUE,name="A2_3"))
  A_beta_3<-omxCbind(A_beta_3,get(paste0("A",i,"_3")), name="A_beta_3")
}


A_mid_3 <-  mxMatrix("Full", nrow=4,ncol=4,
                     labels = NA,
                     values = c(c(rep(0,4)),
                                1,
                                c(rep(0,8)),
                                0.5,sqrt(0.75),
                                0),
                     free = F,
                     name="Amid_3",
                     byrow=TRUE)
A_diag_3 <-  mxMatrix("Full", nrow=4,ncol=4,
                      labels = NA,
                      values = c(c(rep(0,4)),
                                 1,
```

```
                    c(rep(0,8)),
                    0.5,sqrt(0.75),
                    0),
                free = F,
                name="Ad_3",
                byrow=TRUE)
for (i in 1:(nvar3-2)){

  A_zeroh_3 <- mxMatrix("Zero", nrow=4,ncol=4*i,labels=NA, byrow=TRUE,
name="Azh_3")
  A_zerov_3 <- mxMatrix("Zero", nrow=4*i,ncol=4,labels=NA, byrow=TRUE,
name="Azv_3")

  A_mid_3 <- omxRbind(omxCbind(A_mid_3, A_zerov_3),
                omxCbind(A_zeroh_3,A_diag_3),name= "A_mid_3")
}


A_error_3 <- mxMatrix("Full", nrow=nvarall3,ncol=1,labels=NA, free = F,
                values = c(1,
                    rep(0,nvarall3-1)), byrow=TRUE, name="Ae_3")
A_bottom_3 <- mxMatrix("Zero", nrow=1,ncol=4*(nvar3-1),labels=NA, byrow=TRUE,
name="Ab_3")




Am_3 <- omxCbind(A1_3,omxRbind(A_beta_3,
                    A_mid_3,A_bottom_3),A_error_3,name="A",
            dimnames = list(allVariables3,allVariables3))


##S matrix


S1_3 <-  mxMatrix("Zero", nrow=nvarall3,ncol=nvar3,labels=NA, byrow=TRUE,
name="S1_3")
S_top_3 <- mxMatrix("Zero", nrow=nvar3,ncol=4*(nvar3-1),labels=NA, byrow=TRUE,
name="St_3")


S_mid_3 <- mxMatrix("Diag", nrow=4,ncol=4,
               labels=c(paste0("var_",varnames3[2])
                   ,NA,
                   paste0("var_",varnames3[2]),
                   NA),
               values = c(0.2, 0, 0.2, 0),
               free= c(T,F,T,F),
               byrow=TRUE, name="Smid_3")
for (i in 2:(nvar3-1)){
```

```
  S_diag_3 <- mxMatrix("Diag", nrow=4,ncol=4,
                  labels=c(paste0("var_",varnames3[i+1]),NA,
                         paste0("var_",varnames3[i+1]),NA),
                  values = c(0.2, 0, 0.2, 0),
                  free= c(T,F,T,F),
                  byrow=TRUE, name="Sd_3")
  S_covv_3 <-mxMatrix("Zero", nrow=0,ncol=4,labels=NA, byrow=TRUE, name="Sch_3")
  S_covh_3 <-mxMatrix("Zero", nrow=4,ncol=0,labels=NA, byrow=TRUE, name="Sch_3")
  for (j in 2:i){
    S_covh_3 <- omxCbind( S_covh_3 , mxMatrix("Diag", nrow=4,ncol=4,

labels=c(rep(c(paste0("cov_",varnames3[j],"_",varnames3[i+1]),NA),2)),
                                    values = 0,
                                    free= c(T,F,T,F),
                                    byrow=TRUE, name="S_ch_3"))
    S_covv_3 <- omxRbind( S_covv_3 , mxMatrix("Diag", nrow=4,ncol=4,

labels=c(rep(c(paste0("cov_",varnames3[j],"_",varnames3[i+1]),NA),2)),
                                    values = 0,
                                    free= c(T,F,T,F),
                                    byrow=TRUE, name="S_cv_3"))
  }


  S_mid_3 <-  omxRbind(omxCbind(S_mid_3, S_covv_3),
                  omxCbind(S_covh_3, S_diag_3))
}



S_error_3 <- mxMatrix("Full", nrow=nvarall3,ncol=1,
                 values = c(rep(0,nvarall3-1),
                        1),
                 free= c(rep(F,nvarall3-1),T),
                 labels = c(rep(NA,nvarall3-1),"phi2"),
                 byrow=TRUE, name="Se_3")
S_bottom_3 <- mxMatrix("Zero", nrow=1,ncol=4*(nvar3-1),labels=NA, byrow=TRUE,
name="Sb_3")
Sm_3<- omxCbind(S1_3,
             omxRbind(S_top_3,S_mid_3,S_bottom_3),
             S_error_3,
             name="S", dimnames = list(allVariables3,allVariables3))

##F matrix
F1_3 <- mxMatrix("Iden", nrow=nvar3,ncol=nvar3,labels = NA,byrow=TRUE,
name="F1_3")
F2_3 <- mxMatrix("Zero", nrow=nvar3,ncol=nvarall3-nvar3, labels =NA,byrow=TRUE,
name="F2_3")
Fm_3 <-  omxCbind(F1_3,F2_3,name="F", dimnames = list(manifests3,allVariables3))
```

```
##M matrix
Mm_3 <- mxMatrix("Full", nrow=1,ncol=nvarall3,dimnames=list(NULL,allVariables3),
            values= 0,
            free=c(rep(T,nvar3),rep(F,nvarall3-nvar3)),
            labels=c(paste0("mean_",varnames3),rep(NA,nvarall3-nvar3))
            , byrow=TRUE, name="M")
fitFunction_3 <- mxFitFunctionML()
expFunction_3 <- mxExpectationRAM(A ="A",S ="S",F = "F",M ="M",
                        dimnames= allVariables3)

model_3 <- mxModel("SEM_BW_HLA_3",
             type= "RAM",
             Am_3,Sm_3,Fm_3,Mm_3,
             mxData(modelData3, type = "raw"),
             expFunction_3,
             fitFunction_3 )


##combine the 3 submodels together
minus2ll <- mxAlgebra(expression=SEM_BW_HLA.fitfunction +
SEM_BW_HLA_2.fitfunction + SEM_BW_HLA_3.fitfunction  ,
               name="minus2loglikelihood" )
obj <- mxFitFunctionAlgebra("minus2loglikelihood")
model <- mxModel(model="IV Model", model_1, model_2, model_3 ,minus2ll, obj)
result_TH <- mxTryHard(model)

summary(result_TH)

##calculate P values
Zscore_base <- summary(result_TH)$parameters[,5]/summary(result_TH)$parameters[,6]
Pval_base <- 2*(1-pnorm(abs(Zscore_base),0,1))
```
